# Non-canonical TLR signaling restricts cytosolic LPS detection

**DOI:** 10.1101/2025.03.10.642106

**Authors:** Shriram Ramani, Sara Cahill, Matthew Finnegan, Laurel Stine, Abigail Sondrini, Kiera Clayton, Liraz Shmeul-Galia, Fiachra Humphries

## Abstract

Intracellular sensing of lipopolysaccharide (LPS) is an essential component of pathogen detection that governs the innate immune response. However, how this process is controlled to maintain homeostasis and resolve inflammation is unclear. Here, we show that MARCO is a decoy LPS sensor crucial for restraining caspase 11 activity and the non-canonical inflammasome. Remarkably, MARCO expression is controlled by a non-canonical TLR signaling pathway involving the metabolite itaconate, the autophagy adaptor protein p62, and the transcription factor NRF2. In the presence of IFN, non-canonical TLR signaling is impaired and NRF2 dependent gene expression is terminated. Thus, impairing MARCO expression and licensing optimal activation of the non-canonical inflammasome. Loss of MARCO augments non-canonical inflammasome activation and sensitizes mice to septic shock. Together, this study identifies MARCO as a previously unknown LPS sensor that is regulated by a non-canonical TLR signaling pathway and reveals an intricate homeostatic switch that allows for optimal immune responses and resolution of inflammation.

## Introduction

Inflammasomes are signaling complexes formed following the detection of damage associated molecular patterns (DAMPs) (*1, 2*). Nucleotide-binding domain (NBD), leucine-rich repeat (LRR) protein (NLRs) can recruit the adaptor protein apoptosis-associated speck-like protein containing a CARD (ASC) and the protease caspase 1. Active caspase 1 can then induce the proteolytic maturation of IL-1β and IL-18 (*3*). In addition, inflammasome activation can result in an inflammatory form of cell death known as pyroptosis (*4, 5*). Pyroptosis is triggered downstream of inflammasome assembly and facilitates the release of IL-1β and IL-18, via gasdermin (GSDM) pores, and larger DAMPs following plasma membrane rupture (PMR) (*6–8*). During pyroptosis, specific GSDM proteins can be cleaved by inflammatory caspases, such as caspase 1 or 11. Upon cleavage and release, the GSDM N-terminal domain oligomerizes and forms membrane pores, which can lead to the collapse of the membrane and cell death (*4, 9–13*).

Cytosolic lipopolysaccharide (LPS) is sensed by caspase 11 (caspase 4 in humans), which triggers self-cleavage and subsequent cleavage and activation of gasdermin D (GSDMD), and the activation of pyroptosis (*4, 14, 15*). Potassium efflux induced by GSDMD pores indirectly triggers non-canonical NLRP3 oligomerization and the maturation and release of IL-1β and IL-18 (*16*). Bacterial derived LPS can be released into circulation via the complement system, antibiotics, and antimicrobial peptides (*17*). Additionally, free LPS can associate with extracellular vesicles (EVs) *in vivo* to gain access to the cytosol via CD14 (*17, 18*). However, how this is controlled to maintain homeostasis is unknown. In this study, we have identified a previously known branch of the TLR signaling downstream of Myd88 which induces an NRF2-dependent gene program that is restricted by the presence of type I interferon (IFN). In our study we identify and characterize one product of this signaling event, <u>m</u>acrophage-<u>a</u>ssociated <u>r</u>eceptor with <u>co</u>llagenous structure (MARCO) which functions as a decoy LPS receptor. MARCO is a scavenger receptor uniquely expressed on macrophages. MARCO has been implicated in the clearance of pathogens and apoptotic debris, tumorigenesis, and autoimmunity (*19–24*). MARCO contains a scavenger receptor cysteine rich (SRCR) domain, which is required for ligand binding (*22*). However, MARCO contains no signaling domain. Despite these findings, there is a lack of genetic studies and a paucity in our understanding of the functional role that MARCO plays in innate immunity and how MARCO expression is regulated on tissue resident macrophages. Remarkably, we found that interferon (IFN) suppresses MARCO expression on macrophages. Mechanistically, exposure of macrophages to IFN results in impaired NRF2-dependent gene expression and thus reduction of MARCO expression on the cell surface. In the absence of MARCO, the progression of septic shock was exacerbated. IFN-mediated suppression of MARCO expression is a previously unknown innate checkpoint that facilitates the recognition of LPS by caspase 11 and the progression of septic shock.

## Results

### MARCO is an LPS binding protein

To identify previously unknown LPS sensors, we performed a proteomic screen using biotinylated LPS. THP1 derived human macrophages were incubated with biotin-LPS, followed by enrichment of LPS binding proteins using streptavidin beads. Mass spectrometry analysis identified MARCO as an LPS binding protein **(Figure 1A-C)**. While endogenously expressed MARCO has not been identified as a LPS binding protein, *in vitro* studies have shown that soluble MARCO (sMARCO) binds LPS via the SRCR domain, supporting that LPS interacts with MARCO physiologically (*25*). Furthermore, phage display finds that sMARCO preferentially binds hydrophobic peptides via the cystine-rich SRCR domain, suggesting that MARCO could interact with the hydrophobic region of LPS (*26, 27*). Acidic clusters on the SRCR domain are also hypothesized to potentiate electrostatic interactions and mediate ligand binding (*28*). We next evaluated the expression level of MARCO in primary murine bone marrow derived macrophages (BMDMs). However, MARCO was not expressed at the protein level in BMDMs. We thus compared MARCO expression in non-differentiated and differentiated THP1 cells. PMA treatment results in the induction of MARCO in THP1 derived macrophages **(Figure 1D)**. We next evaluated if inflammatory signals could stimulate MARCO expression in BMDMs. The toll like receptor (TLR) 2 ligand Pam3CSK4 induced robust MARCO expression. However, TLR4 and TLR3 agonists, LPS and Poly (I:C), respectively, both failed to induce MARCO expression **(Figure 1E)**. Given that TLR2 signaling does not induce type I IFN signaling via TBK1 and IRF3, we hypothesized that autocrine IFN signaling restricted MARCO expression. Indeed, treatment with the JAK1 inhibitor ruxolitinib restored MARCO expression in LPS and Poly (I:C) treated macrophages. STING activation using diABZI failed to induce MARCO expression, suggesting that TLR signaling is required for MARCO expression on macrophages (**Figure 1F-G**). We next re-evaluated LPS binding to MARCO in LPS primed BMDMs in the presence of ruxolitinib. Ruxolitinib treatment resulted in the precipitation of MARCO by biotin-LPS (**Figure 1H-J**). Biotin-LPS bound to recombinant MARCO by ELISA (**Figure 1K**) and colocalized with MARCO in Pam3CSK4 primed BMDMs (**Figure 1L**). FITC-labelled LPS also co-localized with MARCO **(Figure S1A)**. Thus, MARCO is a previously unknown LPS binding protein controlled by IFN.

**Figure 1:**
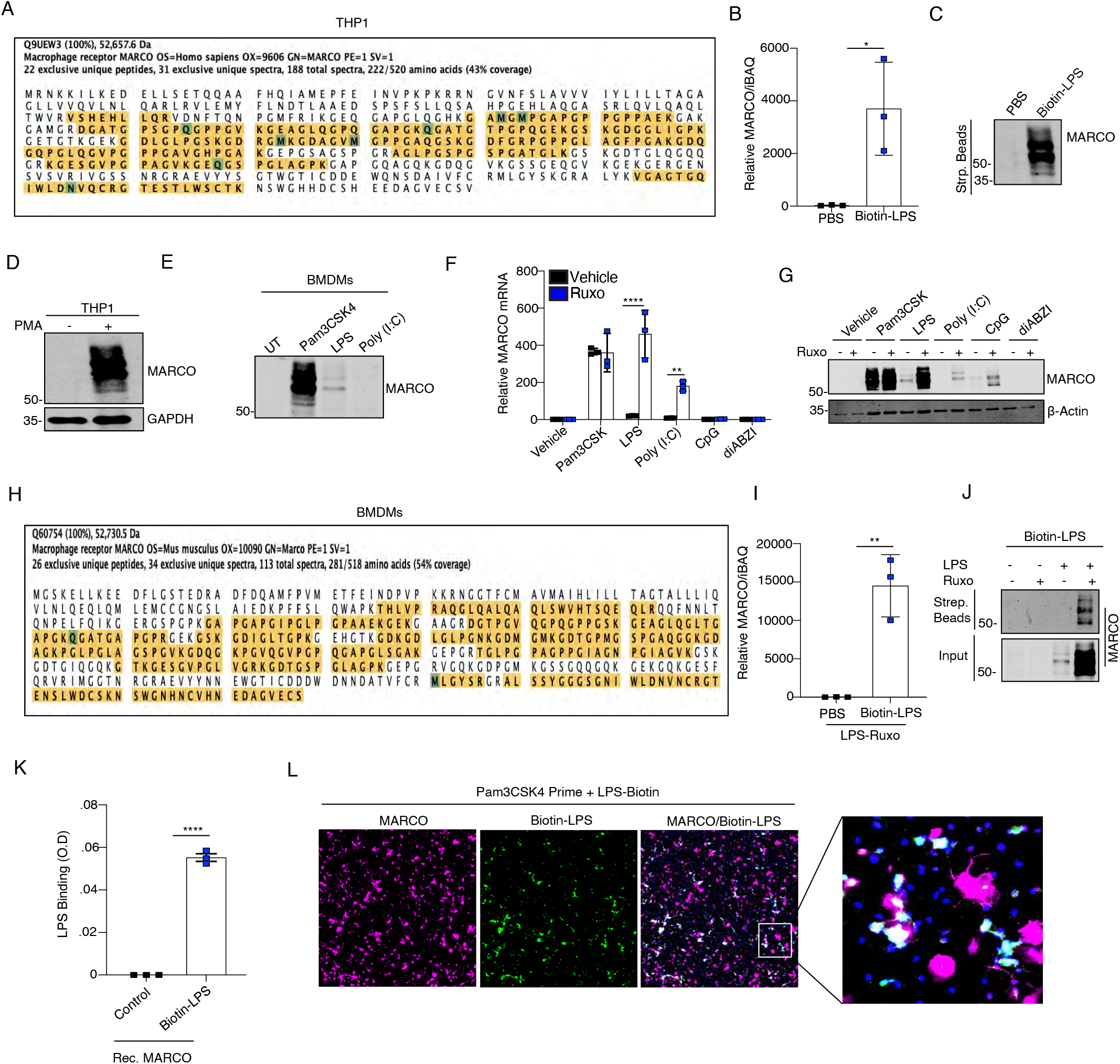
MARCO is an LPS binding protein. **(A-B)** Peptide map (A) and relative abundance (B) of MARCO eluted from streptavidin beads incubated in PMA differentiated THP1 monocytes treated with PBS or biotin-LPS (500ng/mL) for 1 hour. **(C)** Immunoblot analysis of MARCO from streptavidin elution from A. **(D)** Immunoblot analysis of MARCO and GAPDH in THP1 cells treated with or without PMA (50ng/mL) for 48 hours. **(E)** Immunoblot analysis of MARCO and GAPDH in WT BMDMs treated with Pam3CSK4 (1μg/mL), LPS (100ng/mL) or Poly (I:C) 25 μg/mL for 24 hours. **(F)** RT-qPCR analysis of *Marco* mRNA in WT BMDMs treated with Pam3CSK4 (1μg/mL), LPS (100ng/mL), Poly (I:C), CpG DNA or diABZI (500ng/mL) for 24 hours. **(G)** Immunoblot analysis of MARCO and β-Actin in WT BMDMs with the indicated ligands with and without ruxolitinib (1μM) for 24 hours. **(H-I)** Peptide map (H) and relative abundance (I) of MARCO in eluted from streptavidin beads incubated in WT BMDMs pre-treated with ruxolitinib (100nM) and LPS (100ng/mL) for 24 hours followed by treatment with PBS or biotin-LPS (500ng/mL) for 1 hour. **(J)** Immunoblot analysis of MARCO from streptavidin elution and input from H. **(K)** ELISA assay of immobilized recombinant MARCO incubated with biotin LPS. **(L)** Immunofluorescence staining of MARCO (pink), biotin-labelled LPS (green), and Hoechst (blue) in WT BMDMs primed with Pam3CSK4 (1μg/mL) for 24 hours followed by incubation with biotin-LPS (500ng/mL) for 1 hour. B, F, I, K pooled data from 3 independent experiments. A, C-E, G, H, J, L representative images from 3 independent experiments. *P<0.05, **P<0.01, ***P<0.001 student’s t-test.

### MARCO is an IFN-restricted gene

We next assessed the ability of IFN to restrict MARCO expression. Using an unbiased approach, we sequenced RNA isolated from LPS treated WT and *Ifnar^-/-^*BMDMs. Interestingly, MARCO was one of the most upregulated genes in LPS-treated IFNAR-deficient cells when compared to LPS treated WT cells **(Figure 2A-C)**. Expression of the IFN-stimulated gene (ISG), *Cxcl10*, was lost in *Ifnar^-/-^* BMDMs **(Figure 2C**). Sequencing of RNA from LPS and ruxolitinib treated cells also identified MARCO as one of the most upregulated genes in ruxolitinib treated cells **(Figure S1B-E)**. Pre-treatment of cells with anti-IFNAR blocking ^antibody^ **(Figure 2D)**, with ruxolitinib or tofacitinib **(Figure 2E, S2A)**, and treatment of *Irf3^-/-^* or *Ifnar^-/-^* cells **(Figure 2F-G)** all resulted in robust LPS-induced MARCO expression. Indirect stimulation of IFNβ via the addition of exogenous recombinant IFNβ to *Irf3^-/-^* cells restricted augmented MARCO **(Figure S2B)**. We next evaluated the effect of IFN exposure on macrophages expressing MARCO. Exogenous IFNβ treatment suppressed MARCO in Pam3CSK4 primed macrophages **(Figure S2C)**. Flow cytometry analysis demonstrated complete inhibition of TLR2 induced MARCO expression in the presence of type I (IFNβ) or type II (IFNγ) IFN **(Figure 3A-D, S3)**. Addition of exogenous IFNα, β or γ all suppressed mRNA and protein levels of MARCO in Pam3CSK4 primed macrophages (**Figure 3E-F**). Further, exogenous IFNγ suppressed MARCO in both WT and *Ifnar^-/-^*cells (**Figure 3G)**. MARCO is expressed basally in CD14-monocyte derived human macrophages **(Figure 3H)**. Consistently, exposure of primary human macrophages to IFNβ resulted in the complete suppression of MARCO expression **(Figure 3H)**. Further, JAK1 inhibition using ruxolitinib inhibited IFN mediated downregulation of MARCO in primary human macrophages **(Figure 3I)**. Given that MARCO was expressed basally in human macrophages, we assessed the relative expression of MARCO in naïve CD14+ monocytes versus monocytes cultured in M-CSF, GM-CSF, or M-CSF+GM-CSF for 7 days. M-CSF induced MARCO expression in cultured human macrophages **(Figure 3J)**. This is unsurprising given that MARCO is expressed in tissue resident macrophages that are maintained by basally produced M-CSF. Collectively, these data demonstrate that MARCO is an IFN-restricted gene.

**Figure 2:**
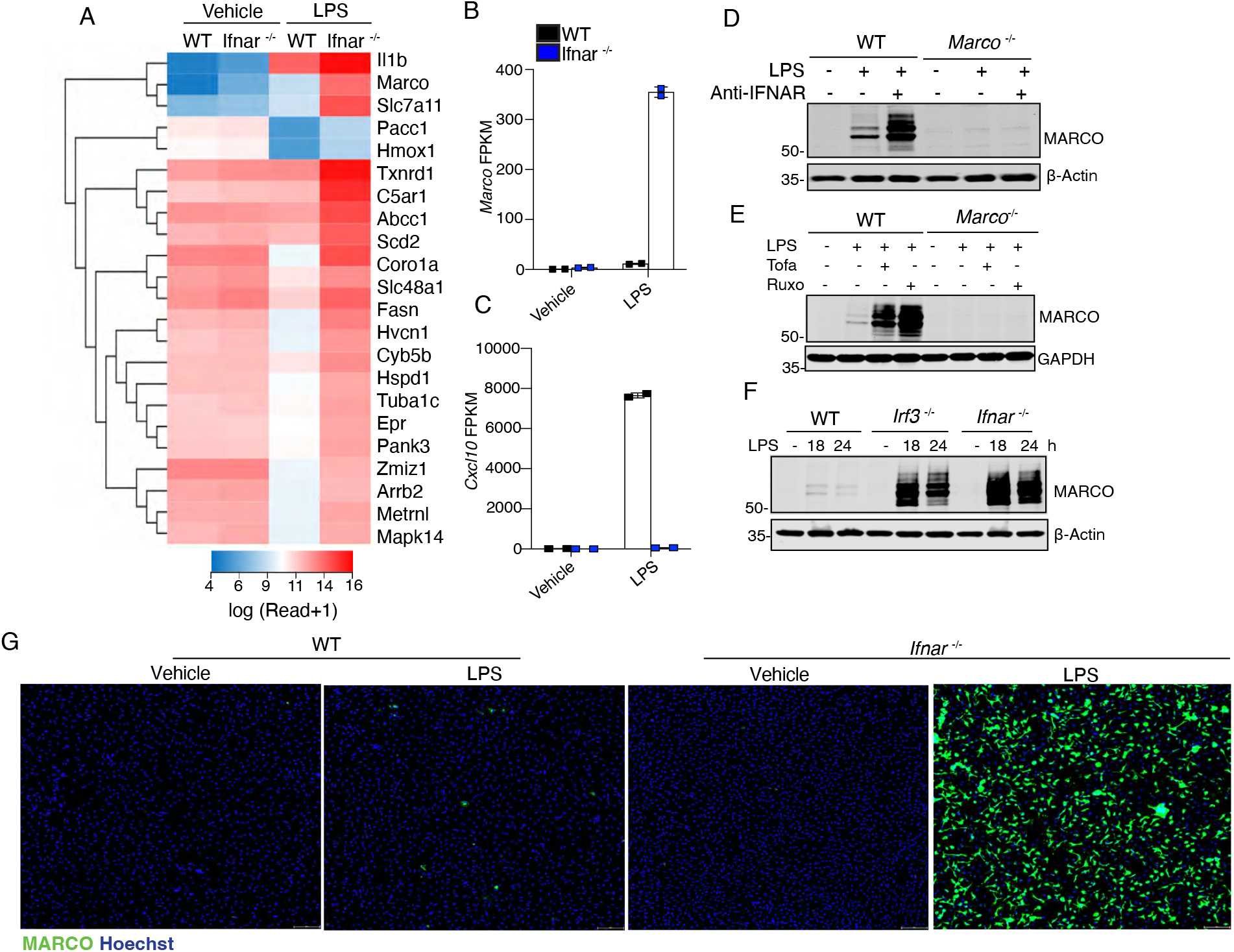
MARCO is an IFN-restricted gene. **(A)** Heat map of differentially expressed genes from RNA-seq analysis of WT and *Ifnar-/-* BMDMs treated with LPS (100ng/mL) for 6 hours. **(B)** Heat map of the top 20 most significantly upregulated genes in LPS (100ng/mL) treated *Ifnar^-/-^*cells relative to WT LPS (100ng/mL) treated cells. (**C-D)** FPKM values of *Marco* (C) and *Cxcl10* (D) from A. **(E)** Immunoblot analysis of MARCO and β-actin in WT BMDMs treated with anti-IFNAR (10ng/mL) for 1 hour, followed by LPS (100ng/mL) for 24 hours. **(F)** Immunoblot analysis of MARCO and GAPDH in WT BMDMs treated with ruxolitinib (1μM) or tofacitinib (1μM) for 1 hour, followed by LPS (100ng/mL) 24 hours. **(G)** Immunoblot analysis of MARCO and β-actin in WT, *Ifnar ^-/^,^-^* and *Irf3-/-* BMDMs treated with LPS (100ng/mL) for 18 and 24 hours. **(H)** Immunofluorescence staining of MARCO (green) and Hoechst (blue) in WT and *Ifnar-/-* BMDMs treated with LPS (100ng/mL) for 24 hours. A-D average of two-independent experiments. E-H representative images from 3 independent experiments

**Figure 3:**
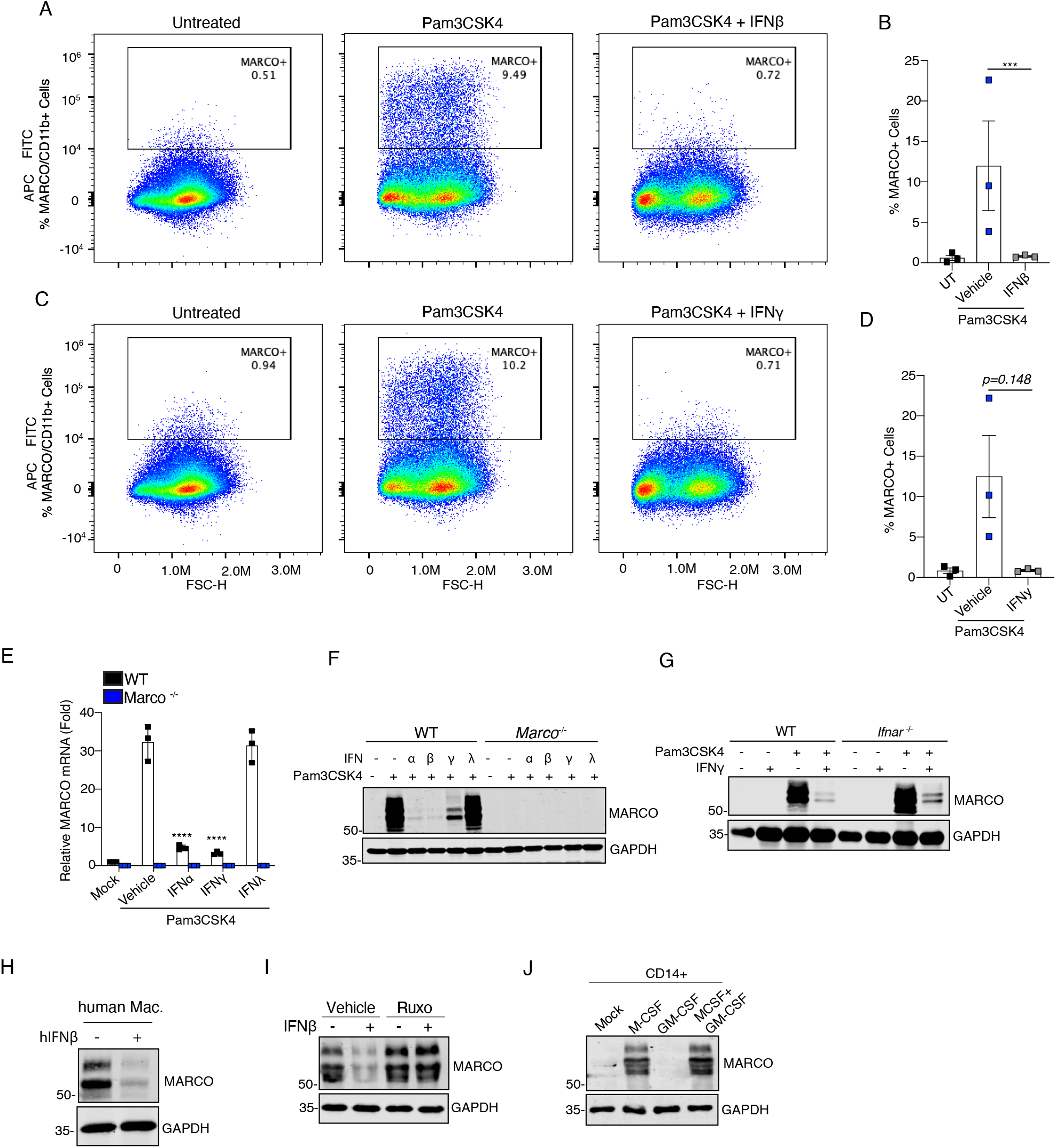
IFN exposure inhibits MARCO in murine and human macrophages. **(A-B)** Representative flow cytometry plots (A) and quantification (B) of CD45+, CD11b+, and MARCO+ cells treated with Pam3CSK4 (1μg/mL) for 1 hour followed by IFNβ (10ng/mL) for 24 hours. **(C-D)** Representative flow cytometry plots (C) and quantification (D) of CD45+, CD11b+, and MARCO+ cells treated with Pam3CSK4 (1μg/mL) for 1 hour followed by IFNγ (10ng/mL) for 24 hours. **(E)** RT-qPCR analysis of *Marco* mRNA in WT BMDMs treated with Pam3CSK4 (1μg/mL) for 1 hour, followed by treatment with IFNα (10ng/mL), IFNγ (10ng/mL), and IFNλ (10ng/mL) for 5 hours. **(F)** Immunoblot analysis of MARCO and GAPDH in WT and *Marco^-/-^*BMDMs treated with Pam3CSK4 (1μg/mL) for 1 hour, followed by treatment with IFNα (10ng/mL), IFNβ (10ng/mL), IFNγ (10ng/mL), and IFNλ (10ng/mL) for 24 hours. **(G)** Immunoblot analysis of MARCO and GAPDH in WT and *Ifnar-/-* BMDMs treated with Pam3CSK4 (1μg/mL) for 1 hour, followed by IFNγ (10ng/mL) for 24 hours. **(H)** Immunoblot of MARCO and GAPDH in lysates from primary human monocyte-derived macrophages treated with hIFNβ (10ng/mL) for 24 hours. **(I)** Immunoblot of MARCO and GAPDH of lysates from primary human monocyte derived macrophages treated with hIFNβ (10 ng/mL) with or without ruxolitinib (1μM) for 24 hours. **(J)** Immunoblot of MARCO and GAPDH in human primary CD14+ monocytes differentiated with, or without, recombinant M-CSF (50ng/mL) and/or GM-CSF (50 ng/mL) for 7 days. B, D-E pooled biological replicates from 3 independent experiments. A, C, F, G are representative images from 3 independent experiments. H-J are representative images from 3 independent donors. **P<0.01, ***P<0.001student’s t-test.

### NRF2 stabilization is restricted by IFN exposure

Given the limited literature on transcription factors required for MARCO expression, we performed a bioinformatic analysis on transcription factor binding sites in open chromatin regions (OCRs) of the MARCO locus. Analysis of ImmGen databases revealed that MARCO expression is limited to the macrophage lineage **(Figure 4A)**. Further, OCR analysis under low and high stringency models identified 4 putative MARCO transcription factors, *JunD, Nfe2l2 (*NRF2*), cFos,* and *Smarcc1* **(Figure 4A-C)**. The nuclear factor erythroid 2-related factor 2 (NRF2) small molecule activators dimethyl fumarate (DMF) and diethyl maleate (DEM) both induced MARCO **(Figure S4A-B)**. Given that the NRF2-dependent gene *Hmox*1 was also elevated in *Ifnar*^-/-^ or ruxolitinib treated cells **(Figure S5A-B),** we evaluated the requirement of NRF2 for MARCO expression. NRF2 is a transcription factor that responds to oxidative stress and induces antioxidant and metabolic gene expression (*29*). Under homeostatic conditions, the E3 ligase kelch-like ECH-associated protein 1 (KEAP1) constitutively degrades NRF2 via K48-linked ubiquitination and proteasomal degradation (*30, 31*). Exposure of macrophages to electrophilic or oxidative stress results in the inhibition of KEAP1 and the stabilization of NRF2 (*30, 32*). Deletion of NRF2 (*Nfe2l2^-/-^*) resulted in the complete loss of MARCO at a transcriptional and protein level **(Figure 4D-G, Figure S5C)**. The NRF2 kinase ERK was also required for MARCO expression **(Figure S5D-E)**. WT and *Nfe2l2*^-/-^ BMDMs displayed comparable levels of LPS induced IFN **(Figure S5F-G)** and expression of the ISGs *Cxcl10* and *Ifit1* **(Figure S5H-I)**. Thus, NRF2 is required for MARCO expression. Given that IFN exposure potently suppresses MARCO expression, we next evaluated the effect of IFN exposure on NRF2 stabilization. Remarkably, we failed to observe stabilization of NRF2 in LPS treated WT BMDMs **(Figure 4H)**. However, IFNAR-deficient BMDMs displayed robust stabilization and nuclear localization of NRF2 **(Figure 4H-I)**. In contrast to other studies, this suggests that autocrine IFN produced downstream of LPS stimulation creates a barrier that limits NRF2 dependent gene expression in macrophages and primes macrophages towards a pro-inflammatory state. TLR2-stimulation, which does not induce any IFN, induced a significant increase in NRF2 stabilization **(Figure 4J)**. Furthermore, exogenous IFNβ treatment impaired Pam3CSK4 induced NRF2 stabilization **(Figure 4K-L)**. PMA stimulation also induced *MARCO* and the other NRF2 dependent genes *HMOX1* and *SLC7A11* in human THP1 monocytes **(Figure 4M-O)**. M-CSF stimulation also induced *MARCO* and the other NRF2 dependent genes *HMOX1* and *SLC7A11* in human PBMC derived CD14+ monocytes **(Figure 4P-R)**.

**Figure 4:**
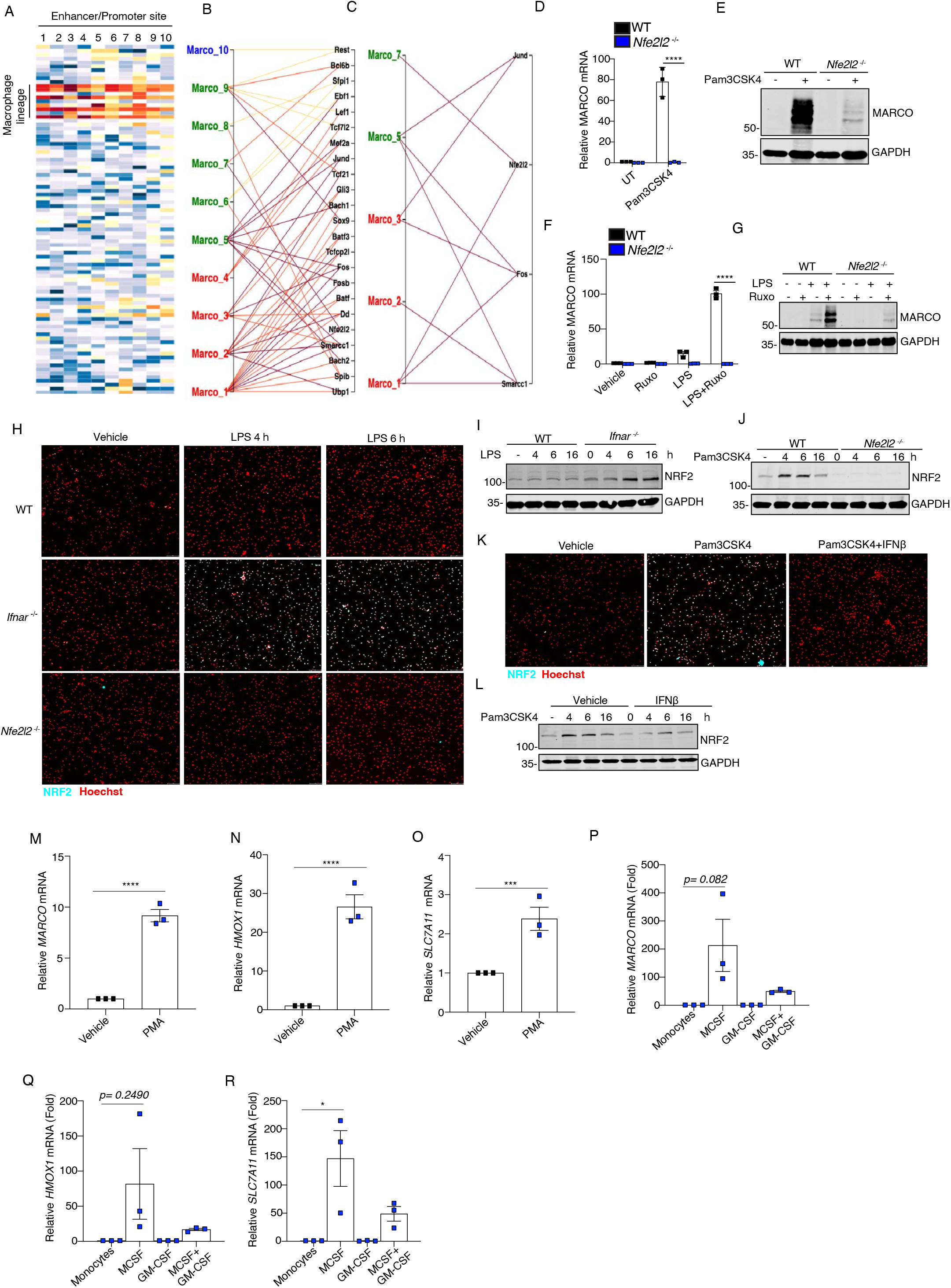
IFN suppresses NRF2 stabilization. **(A-C)** Open chromatin region binding sites in the *Marco* locus. *Data from ImmGen*. **(D)** RT-qPCR analysis of *Marco* mRNA in WT and *Nfe2l2-/-*BMDMs treated with Pam3CSK4 (1μg/mL) for 6 hours. **(E)** Immunoblot analysis of MARCO and GAPDH in WT and *Nfe2l2-/-* BMDMs treated with Pam3CSK4 (1μg/mL), for 24 hours. **(F)** RT-qPCR analysis of *Marco* mRNA in WT and *Nfe2l2-/-* BMDMs treated with ruxolitinib (1μM) for 1 hour, followed by LPS (100ng/mL) for 5 hours. **(G)** Immunoblot analysis of MARCO and GAPDH in WT and *Nfe2l2-/-* BMDMs treated with ruxolitinib (1μM) for 1 hour, followed by LPS (100ng/mL) for 24 hours. **(H)** Immunofluorescence staining of NRF2 (cyan) and Hoechst (red) in WT, *Ifnar-/-,* and *Nfe2l2-/-* BMDMs treated with LPS (1μM) for 4 and 6 hours. **(I)** Immunoblot analysis of NRF2 and GAPDH in WT and *Ifnar-/-* BMDMs treated with LPS (100ng/mL) for the indicated times. **(J)** Immunoblot analysis of NRF2 and GAPDH in WT and *Nfe2l2-/-* BMDMs treated with Pam3CSK4 (1μg/mL) for the indicated times. **(K)** Immunofluorescence staining of NRF2 (cyan) and Hoechst (red) in WT BMDMs treated with Pam3CSK4 (1μg/mL) for 1 hour, followed by treatment with or without IFNβ (10ng/mL) for 5 hours. **(L)** Immunoblot analysis of NRF2 and GAPDH in WT BMDMs pre-treated Pam3CSK4 (1μg/mL) with followed by treatment with IFNβ (10ng/mL) for the indicated times. **(M-O)** RT-qPCR of THP-1s with and without PMA (50 ng/mL) for *Marco* (M), *Hmox1* (N), and *Slc7a11* (O) mRNA. **(P-R)** RT-qPCR of primary CD14+ monocytes differentiated with recombinant M-CSF (50ng/mL) and/or GM-CSF (50 ng/mL) for 7 days for *Marco* (P), *Hmox1* (Q), and *Slc7a11* (R). D, F, M-R pooled biological replicates from 3 independent experiments. A-C data from *Immgen*. E, G, H-L representative images from 3 independent experiments. **P<0.01, ***P<0.001, ****P<0.0001 student’s t-test.

### ROS and p62 aggregation are required for NRF2 stabilization and MARCO expression

We next evaluated the mechanism by which NRF2 is stabilized in the absence of IFN. Previous studies have demonstrated that metabolic changes downstream of TLR4 activation result in the expression of the enzyme immune responsive gene 1 (IRG1; *Acod1*), which synthesizes the immunomodulatory metabolite itaconate (*33*). Itaconate and its derivative 4-octyl itaconate (4-OI) have been shown to promote NRF2 stabilization via different mechanisms (*34*). Indeed, 4-OI can directly alkylate KEAP1 to inhibit NRF2 degradation and NLRP3 to limit the NLRP3 inflammasome (*30, 35*). Endogenous itaconate can also induce ROS production via the inhibition of succinate dehydrogenase (SDH)(*34, 36, 37*). Itaconate can also modify GSDMD via itaconation to mediate macrophage tolerance (*38*). A recent study has also indicated that itaconate can target the anti-oxidative stress protein peroxiredoxin 5 (*39*). Nitric oxide (NO) has also been implicated in the stabilization of NRF2 via the inhibition of KEAP1(*40*). Using unbiased metabolomics, we profiled the metabolome in Pam3CSK4 treated BMDMs. Itaconate was one of the most abundant metabolites induced following Pam3CSK4 stimulation **(Figure 5A-B)**. Pam3CSK4 stimulation induced robust expression of *Irg1* **(Figure 5C-D)**. Further, TLR2-induced MARCO expression was decreased in IRG1-deficient BMDMs **(Figure 5E-F)**. MARCO expression was also reduced in IRG1-deficient BMDMs treated with LPS and ruxolitinib, suggesting a role for IRG1 and itaconate in the regulation of MARCO expression **(Figure 5G-I)**. Although previous studies have indicated that *Irg1* is an ISG (*41*), *Irg1* expression was comparable between WT and *Ifnar*^-/-^ cells **(Figure S6A)**. Expression of IRG1 was also independent of NRF2 **(Figure S6B)**. NO is synthesized by the ISG iNOS (*Nos2*). Indeed, *Nos2* gene expression was completely lost in *Ifnar^-/-^* cells **(Figure S6C)**. LPS induced NO synthesis in a *Nos2*-dependent manner, while TLR2 stimulation did not result in NO synthesis **(Figure S6D)**. In line with previous studies, NO synthesis was elevated in *Irg1^-/-^* cells (*42*) (**Figure S6E**). Treatment with the NO inhibitor SEIT suppressed LPS induced NO synthesis in WT and *Irg1^-/-^* BMDMs (**Figure S6F-G)**. Ruxolitinib treatment inhibited LPS induced NO synthesis in WT and *Irg1^-/-^*BMDMs (**Figure S6H-I)** due to suppression of the ISG iNOS **(Figure S6J)**. However, SEIT treatment did not impair TLR2-induced MARCO expression (**Figure S6K-L)**. Although IRG1-deficiency resulted in a reduction in MARCO, expression was not completely lost. We thus evaluated additional mechanisms of NRF2 activation. Both ROS and the autophagy adaptor protein, p62, have been implicated in NRF2 activation. Indeed, p62 (*Sqstm1*) has been shown to sequester KEAP1 and allow for stabilization of NRF2 (*43*). However, no direct evidence for p62 mediated NRF2 activation downstream of TLR signaling has been established. Pre-treatment of cells with the ROS scavenger mitoTEMPO suppressed Pam3CSK4 induced NRF2 stabilization (**Figure 5J-K)**. Further, deletion of p62 resulted in the loss of NRF2 stabilization in response to Pam3CSK4 (**Figure 5L-M**). Thus, both ROS and p62 were required for NRF2 stabilization. We next evaluated p62 aggregation in response to Pam3CSK4. Pam3CSK4 induced p62 aggregation in a ROS dependent manner (**Figure 5N-O**). Further, MARCO expression was dependent on ROS and p62 **(Figure 6P-R)**. Additionally, p62 aggregation in response to Pam3CSK4 was found to be partially reduced in *Irg1^-/-^* BMDMs (**Figure S6M-N).** These data demonstrate that MARCO expression and NRF2 activation can occur in an NO-independent and IRG1/p62/ROS dependent manner. Thus, macrophages can only stabilize NRF2 in the absence of IFN through a NO-redundant mechanism. This process is then impaired when the macrophage is exposed to IFN.

**Figure 5:**
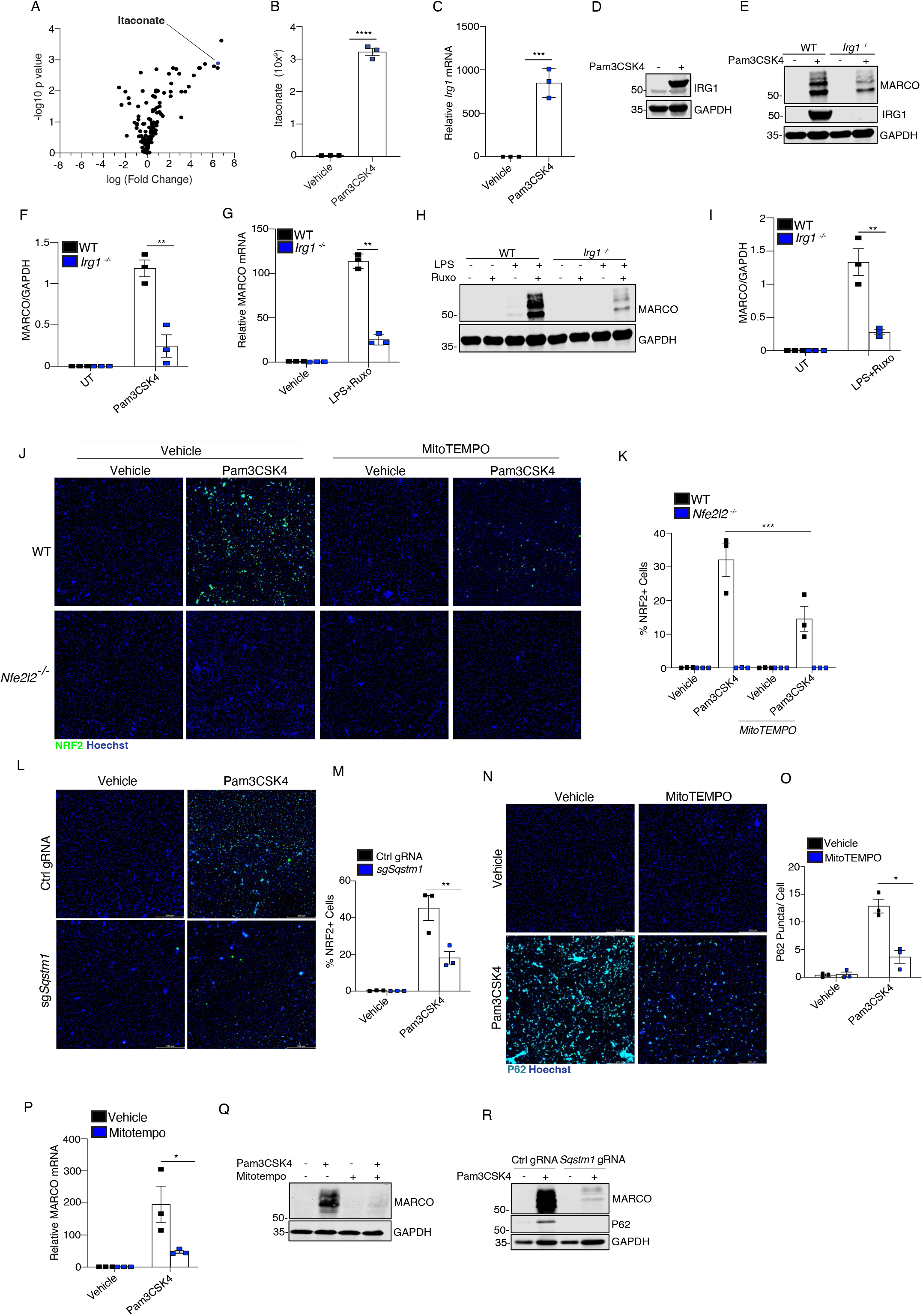
P62 aggregation and ROS are required for NRF2 activity downstream of TLR2. **(A)** Volcano plot of metabolomics analysis of WT BMDMs treated with or without Pam3CSK4 (1μg/mL) for 24 hours. **(B)** Relative abundance of itaconate from A. **(C)** RT-qPCR analysis of *Irg1* mRNA in WT BMDMs treated with or without Pam3CSK4 (1μg/mL) for 6 hours. **(D)** Immunoblot analysis of IRG1 and GAPDH in WT BMDMs treated with or without Pam3CSK4 (1μg/mL) for 24 hours. **(E)** Immunoblot analysis of MARCO, IRG1, and GAPDH in WT and *Irg1^-/-^* BMDMs treated with or without Pam3CSK4 (1μg/mL) for 24 hours. **(F)** Densitometry analysis of MARCO normalized to GAPDH from E. **(G)** RT-qPCR analysis of *Marco* mRNA in WT and *Irg1^-/-^* BMDMs treated with or without ruxolitinib (1μM) for 1 hour followed by LPS (100ng/mL) for 5 hours. **(H)** Immunoblot analysis of MARCO and GAPDH in WT and *Irg1-/-* BMDMs treated with or without ruxolitinib (1μM) for 1 hour followed by LPS (100ng/mL) for 24 hours. **(I)** Densitometry analysis of MARCO normalized to GAPDH from H. **(J-K)** Representative images (J) and quantification (K) of immunofluorescence analysis of NRF2 (green) and Hoechst (blue) on WT and *Nfe2l2^-/-^* BMDMs treated with or without Pam3CSK4 (1μg/mL) and MitoTEMPO (250μM) 6 hours. **(L-M)** Representative images (L) and quantification (M) of immunofluorescence analysis of NRF2 (green) and Hoechst (blue) of BMDMs electroporated with Cas9 and control or sg*Sqstm1* gRNA treated with or without Pam3CSK4 (1μg/mL) for 6 hours. **(N)** Immunofluorescence analysis of p62 (cyan) and Hoechst (blue) of WT BMDMs treated with or without Pam3CSK4 (1μg/mL) and mitoTEMPO (250μM) for 24 hours. **(O)** Quantification of the number of p62 puncta per cell (measured by Hoechst) in (N). **(P)** RT-qPCR of *Marco* mRNA of WT BMDMs treated with or without Pam3CSK4 (1μg/mL) and mitoTEMPO (250μM) for 24 hours. **(Q)** Immunoblot analysis of MARCO and GAPDH expression of WT BMDMs treated with or without Pam3CSK4 (1μg/mL) and MitoTEMPO (250μM) for 24 hours. **(R)** Immunoblot analysis of MARCO, p62, and GAPDH of WT BMDMs electroporated with Cas9 and control or *Sqstm1* gRNA and treated with Pam3CSK4 (1μg/mL) for 24 hours. B, C, F-G, I, K, M, O, and P are pooled biological replicates from 3 independent experiments. A, D-E, H, J, L, N, Q, and R are representative images from 3 independent experiments. *P<0.05, **P<0.01, P<0.001, ****P<0.0001 student’s t-test.

**Figure 6:**
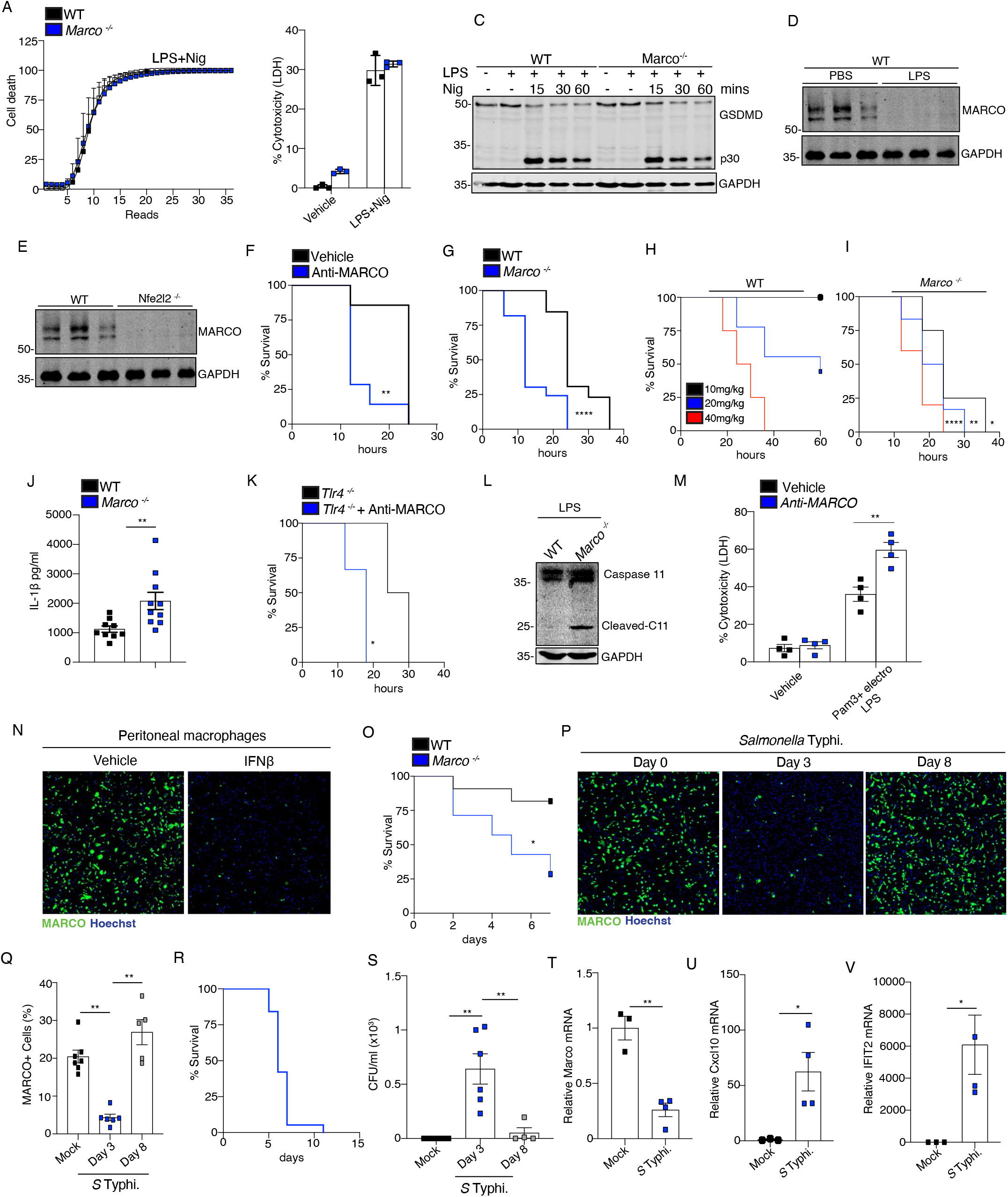
IFNs prime the non-canonical inflammasome by suppressing MARCO. **(A)** Kinetic quantification of SYTOX orange positive cells from WT and *Marco-/-* BMDMs treated with LPS (100ng/mL) for 3 hours and nigericin (10μM) for 1 hour. **(B)** LDH assay on supernatants from WT and *Marco-/-* BMDMs treated with LPS (100ng/mL) for 3 hours and nigericin for 1 hour. **(C)** Immunoblot analysis of GSDMD and GAPDH in lysates from WT and *Marco-/-* BMDMs treated with LPS (100ng/mL) for 3 hours and nigericin (10μM) for the indicated times. **(D)** Immunoblot of MARCO and GAPDH on lysates from digested spleens from mice injected IP with PBS or LPS (10mg/kg) for 6 hours. **(E)** Immunoblot of MARCO and GAPDH on lysates from digested spleens from WT and *Nfe2l2-/-* mice. **(F)** Survival analysis of WT mice administered anti-MARCO antibody (50mg/kg) for 1 hour followed by LPS (40mg/kg). **(G)** Survival analysis of WT and *Marco-/-* mice administered LPS (40mg/kg). **(H-I)** Survival analysis of WT (H) and *Marco-/-* (I) mice administered the indicated concentrations of LPS. **(J)** ELISA analysis of IL-1β in serum from WT and *Marco-/-* mice administered LPS (10mg/kg) for 6 hours. **(K)** Survival analysis of *Tlr4*^-/-^mice administered LPS (40mg/kg) with or without anti-MARCO antibody (20mg/kg). **(L)** Immunoblot of Caspase 11 and GAPDH on lysates from digested spleens from WT and *Marco-/-*mice administered LPS (10mg/kg) for 5 hours. (M) LDH assay on supernatants from primary human monocyte derived macrophages pretreated with Pam3 (1μg/mL) for 6 hours and transfected with LPS (1μg) for 18 hours, with and without the pretreatment of Anti-MARCO blocking antibody (40μg/mL). **(N)** Immunofluorescence of MARCO (green) and Hoechst (blue) on peritoneal macrophages treated with IFNβ (50ng/mL) for 24 hours. **(O)** Survival analysis of WT and *Marco-/-* mice administered 200 μl cecal slurry. **(P-Q)** Representative image (P) and quantification (Q) of immunofluorescence of MARCO (green) and Hoechst (blue) on peritoneal macrophages from mice infected with *Salmonella* Typhi for the indicated times. **(R)** Survival analysis of WT mice infected with *Salmonella* Typhi. **(S)** CFU count on peritoneal lavage fluid from (P). **(T-V)** RT-qPCR of *Marco* (T)*, Cxcl10* (U), and *Ifit2* (V) mRNA from peritoneal macrophages of mice infected with *Salmonella* Typhi for 3 days. A-B, M pooled biological replicates from 3 independent experiments or C, representative images from 3 independent experiments. Representative images from D-E, L, N *n*=3 mice per group. F-K, O, P-V *n*=3-10 mice per group. *P<0.05**P<0.01,****P<0.0001 student’s t-test or Mantel-cox survival analysis.

### MARCO negatively regulates non-canonical inflammasome activation and septic shock

The potent effect IFN has on NRF2 suggests a priming event is required for optimal immune responses and silencing of the anti-inflammatory function of NRF2. We next evaluated the functional significance of MARCO-LPS binding and explored how IFN exposure can counteract this. WT and *Marco^-/-^*BMDMs displayed comparable inflammasome priming (signal 1) with no changes to TLR4 induced gene expression or to the activation of TBK1 and IRF3 **(Figure S7A-F)**. These data are consistent with LPS priming suppressing MARCO expression in BMDMs. As inflammasome priming was unaffected, we next evaluated changes to canonical NLRP3 inflammasome activation. WT and *Marco*^-/-^ BMDMs displayed comparable levels of LPS + nigericin induced pyroptosis, as measured through plasma membrane permeability, LDH release, and GSDMD cleavage **(Figure S9A, 6A-C)**. Cytosolic LPS sensing drives activation of the non-canonical inflammasome, which is dependent on caspase 11 and GSDMD, and contributes to the development of endotoxic septic shock (*4, 16*). Thus, we assessed the role of MARCO expression in non-canonical inflammasome responses in tissue resident macrophages by administering LPS to mice. MARCO is basally expressed on splenic macrophages **(Figure S8A)**. Intraperitoneal delivery of LPS resulted in the loss of MARCO on splenocytes **(Figure 6D)**. Splenic expression of MARCO was dependent on NRF2 **(Figure 6E)**. WT and MARCO-deficient mice displayed a comparable immune cell profile in the spleen **(Figure S8B-L)**. Pre-treatment of mice with an anti-MARCO blocking antibody result in enhanced LPS-induced septic shock **(Figure 6F)**. *Marco*^-/-^mice also displayed enhanced sensitivity to LPS-shock and exhibited an increase in the serum levels of IL-1β **(Figure 6G-J)**. Neutralization of MARCO also enhanced LPS sensitivity in *Tlr4^-/-^* mice **(Figure 6K)**. *Nfe2l2^-/-^*, *Irg1^-/-^,* or mice administered the KEAP1 activator VVD130037 all exhibited enhanced lethality in response to high dose LPS in comparison to WT mice **(Figure S9B-D)**. In line with previous studies (*44*), *Ifnar*^-/-^ mice were protected against LPS-shock **(Figure S9E)**. We next evaluated caspase 11 activation. MARCO-deficient mice displayed enhanced caspase 11 activation in splenocytes when compared to WT mice **(Figure 6L)**. MARCO-blockade using a neutralizing antibody also resulted in enhanced caspase 11 activation in primary human monocyte derived macrophages, as measured by increased LDH release following LPS transfection **(Figure 6M)** Caspase 11 is an ISG and is upregulated in macrophages following IFN exposure as a priming event that precedes non-canonical inflammasome activation (*45*). Thus, IFN induced caspase 11 upregulation is accompanied by the concomitant downregulation of MARCO. Indeed, MARCO-expressing tissue resident macrophages downregulate MARCO following exposure to IFNβ **(Figure 6N)**. Although high dose LPS is useful in studying septic shock responses, it does not fully recapitulate the complexity of sepsis. Polymicrobial sepsis can be induced in mice via the intraperitoneal (IP) administration of cecal slurry. Cecal slurry via IP mimics the symptoms of sepsis without using invasive surgical procedures, such as cecal ligation puncture (CLP). *Marco^-/-^*mice subjected to polymicrobial sepsis displayed enhanced lethality when compared to WT mice **(Figure 6O),** suggesting the protective role of MARCO during physiological sepsis. To dynamically evaluate MARCO expression changes throughout the course of infection, we utilized the LPS bearing bacteria *Salmonella* Typhmurium. *Salmonella* infection resulted in a significant downregulation of MARCO on peritoneal macrophages during the acute phase. Further, MARCO expression was restored during the resolution phase of inflammation **(Figure 6P-R)**. MARCO expression on peritoneal macrophages was inversely correlated with bacterial burden **(Figure 6S)**. These data indicate that MARCO serves as a homeostatic receptor dynamically regulated during pathogen exposure. In addition, loss of MARCO expression correlated with the induction of an ISG signature on peritoneal cells **(Figure 6T-V)**. Together, our data demonstrates that IFN-suppresses MARCO expression on tissue resident macrophages to potentiate the detection of cytosolic LPS and activation of the non-canonical inflammasome **(Figure 7)**.

**Fig 7:**
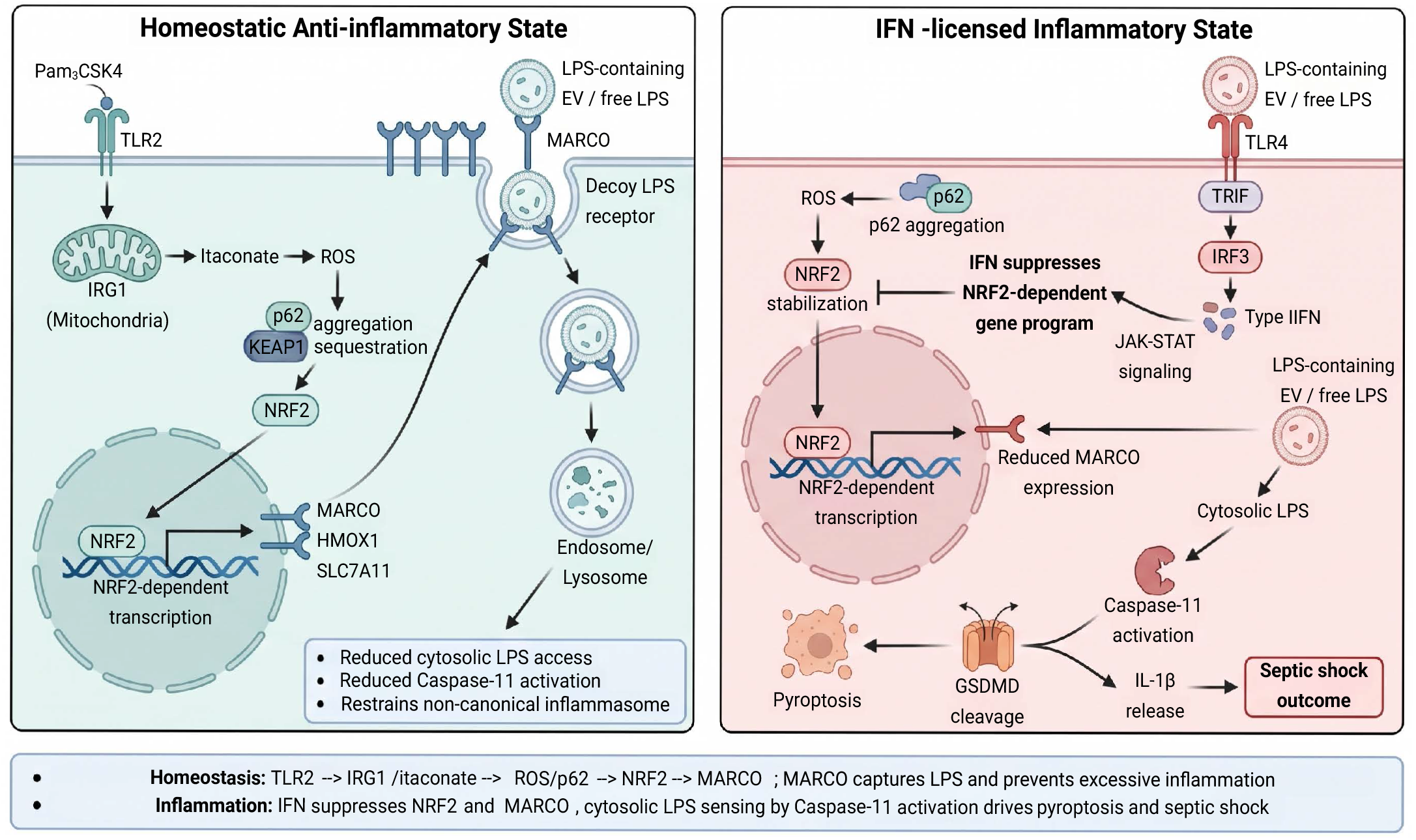
IFN restricts non-canonical TLR signaling and primes cytosolic LPS detection.

## Discussion

Detection of cytosolic LPS and the activation of the non-canonical inflammasome is a key driver of inflammation and septic shock (*16*). Although the signaling events that mediate this process have been identified, there is a lack of understanding of the homeostatic mechanisms that can restrict septic shock. In this study, we have identified a previously unknown non-canonical TLR signaling pathway and feedback innate checkpoint that governs the detection of cytosolic LPS and ensuing activation of the non-canonical inflammasome. Exposure of macrophages to IFN results in the suppression of p62 aggregation and NRF2 stabilization and subsequent downregulation of MARCO in tissue resident macrophages, or suppresses the upregulation of MARCO in monocyte derived macrophages, to maximize the detection of cytosolic LPS and activation of the non-canonical inflammasome. Further, during the resolution phase of an immune response, MARCO is restored to tissue resident macrophages to reestablish tissue homeostasis. Our studies suggest that MARCO can bind directly to LPS and negatively regulate activation of the non-canonical inflammasome. Recent studies have demonstrated that LPS can gain entry to tissue resident macrophages via association with extracellular vesicles (EVs). EVs are then bound by CD14 and internalized and released intracellularly via endosomal damage (*17, 18*). Because MARCO lacks any signaling domain and can also bind EV lipids, it is likely that MARCO can also engage LPS bound EVs and efferocytose and degrade these complexes (*22*). Indeed, *Marco*-deficient mice are highly susceptible to LPS mediated septic shock.

Currently, the exact effector ISG induced by IFN that negatively regulates NRF2 is unknown. An ISG induced following IFN exposure can likely engage the NRF2-KEAP1 complex to impair NRF2 stabilization and suppress the anti-inflammatory function of NRF2. Our data suggests that the NRF2 stabilization downstream of TLR2 or TLR4 is distinct due to the presence of autocrine IFN downstream of TLR4 activation. Indeed, we show that ROS, endogenous itaconate and p62 aggregation are required for NRF2 stabilization in the absence of IFN. Our study also shows, for the first time, that NRF2 stabilization can occur in the absence of NO, but with a requirement for itaconate, ROS, and p62 aggregation. These data explain why NRF2 stabilization is not observed in WT macrophages in response to LPS. Thus, the non-canonical TLR signaling pathway and NRF2 stabilization only occur in macrophages in the absence of IFNs.

Sepsis is a deadly disease that kills over 270,000 people in the United States annually (*46*). Currently, there are no known cures for sepsis (*47*). Our study identifies MARCO as an LPS binding protein that is downregulated by exposure to IFN. Thus, the development of a therapeutic strategy that maintains MARCO levels on tissue resident cells during sepsis may provide benefit for sepsis patients. Indeed, given the safety and efficacy of JAK inhibitors and IFNAR blocking antibodies, it will be of great interest to explore these therapies in sepsis in the clinic.

In summary, our study uncovers the first known metabolic LPS sensor and a previously unknown non-canonical TLR signaling pathway that controls NRF2 activity in macrophages. Further understanding the role MARCO plays in restricting LPS sensing and the signaling events that govern its expression is likely to yield new insights into the pathogenesis and treatment of sepsis and other inflammatory diseases.

## Methods

### Mice

*Marco^-/-^* mice were generated by Taconic using CRISPR. *Irg1^-/-^, Nfe2l2^-/-^, Nos2^-/-^,* and *Tlr4^-/^* mice were obtained from Jackson laboratories. *Ifnar^-/-^* and *Irf3*-/- mice were provided by Prof. Kate Fitzgerald, UMass Chan Medical School. All animal experiments were approved by the Institutional Animal Care Use Committees at the University of Massachusetts Medical School. Animals were kept in a SPF environment.

### Immunoblotting and Immunoprecipitation

Primary BMDMs from mice were cultured in 12-well plates (1×10^6^ cells per mL; 1 mL) or 10-cm dishes (2×10^6^ cells per mL; 10mL). For cell lysate analysis cells were lysed directly in 1X Laemmli sample buffer (BioRad) containing β-Mercaptoethanol (Sigma). For immunoprecipitation of MARCO, cells were treated as indicated, and then collected in 500 µl RIPA buffer, followed by incubation for 15 min on ice. Lysates were incubated with MARCO antibody (human: R&D, murine: Abcam), and protein A–protein G-agarose (Santa Cruz) was incubated with the sample overnight at 4°C. Immunoprecipitates were collected by centrifugation and washed with RIPA buffer. Immunoprecipitates were eluted from beads using 1X Laemmli sample buffer. Samples were resolved by SDS-PAGE, transferred to nitrocellulose membranes (Sigma), and analyzed by immunoblot. Samples were immunoblotted with the indicated antibodies. Proteins were detected using fluorophore-conjugated anti-rabbit (LICOR Biosciences) or anti-sheep secondary antibody (Fisher). Immunoreactivity was visualized using the Odyssey Imaging System (LICOR Biosciences) and densitometry analysis was done using ImageStudio software.

### RT-qPCR

Cells were lysed in TRIzol reagent (Invitrogen), and total RNA was isolated using the Direct-zol RNA miniprep kit (Zymo Research). RNA was quantified by a Nanodrop ND-1000 spectrophotometer (Thermo Scientific), and 1μg of RNA was reverse transcribed using iScript Reverse Transcription Supermix (Bio Rad). qPCR analysis was done on 20 ng of cDNA using the iTaq Universal SYBR Green super-mix reagent (Bio Rad). Murine and human gene expression was normalized to TATA-binding protein (TBP) expression and β-actin, respectively. Relative mRNA expression was calculated by the change in cycling threshold method as 2^-ΔΔC(t)^. Amplification specificity was assessed via melting curve analysis.

Primer sequences are as follows:

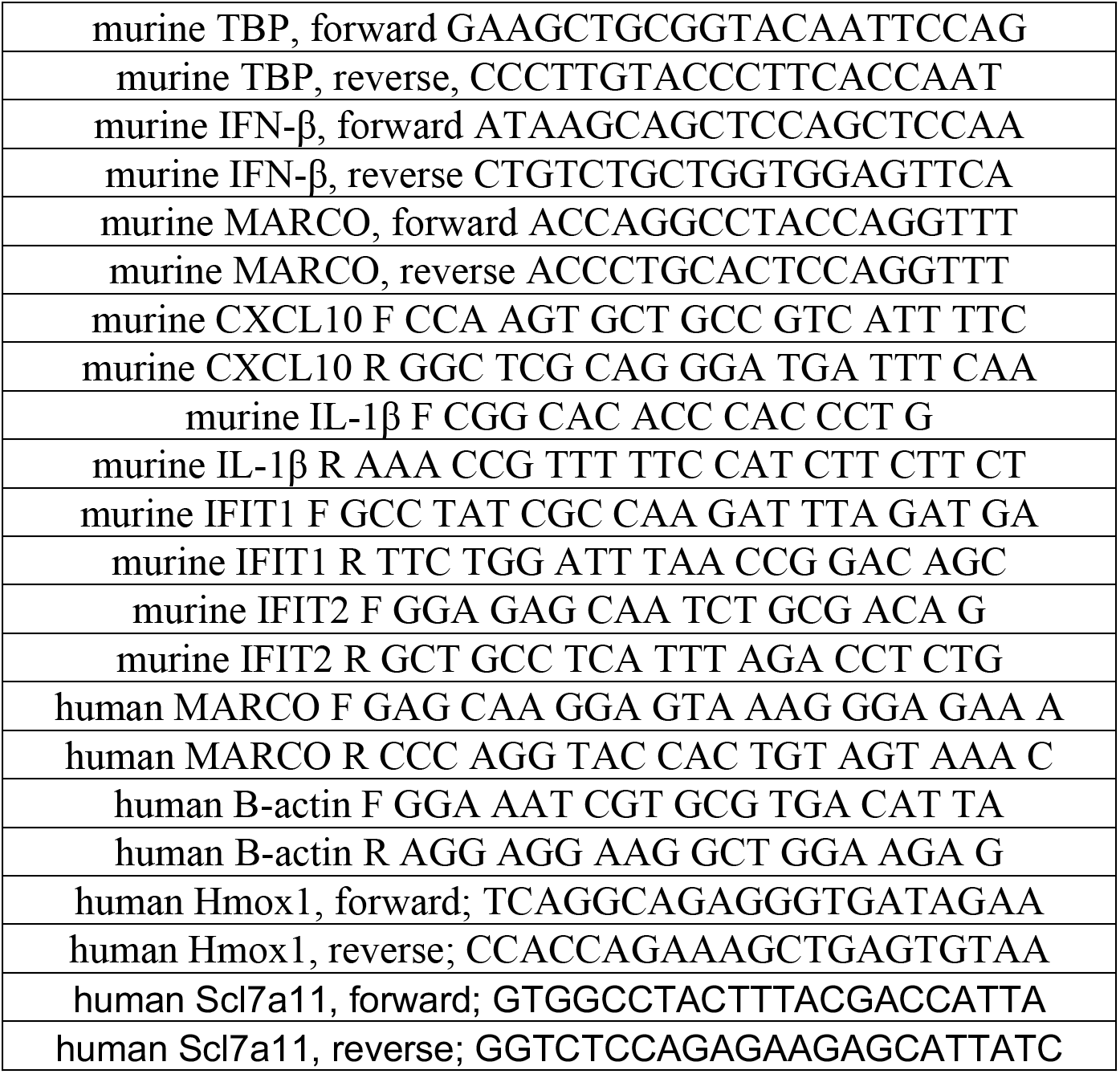

### Kinetic cell death analysis

BMDMs were seeded (0.1 x 10^6^ cells per well) in RPMI-1640 media (ATCC) supplemented with 10% FBS, 1% penicillin and streptomycin and 5% L929 conditioned media in a 96 well plate and rested overnight. To activate the canonical inflammasome, cells were primed with 100 ng/mL LPS (Sigma) for 3 hours and stimulated with 5 μM nigericin (Cayman Chemical). To activate the noncanonical inflammasome, cells were primed with 1 μg/mL Pam3CSK4 (Invitrogen) for 8 hours and transfected with 1 μg LPS (Sigma) using Lipofectamine 3000 (Invitrogen). Cells were stained with 0.25 μM SYTOX orange (Thermo Scientific) and 1 μg/mL Hoechst (Thermo Scientific) and visualized using the BioTek Lionheart FX Microscope (Agilent). The ratio of SYTOX positive cells to Hoechst positive cells were quantified using the BioTek Gen5 Software.

### L929 conditioned media

L929 cells were seeded in T175 flask in DMEM supplemented with 10% FBS, 1% penicillin and streptomycin. Cells were rested for 10 days after reaching confluency at 37 °C in a humidified atmosphere of 5% CO_2_. Conditioned media was passed through a 0.2 μm filter before use.

### ELISA

Measurement of IL-1β was performed using the murine IL-1 ELISA duo set (R&D) according to the manufacturer’s recommended procedure. Measurement of IFN-β was performed using the mouse IFN-beta DuoSet (Biotechne) per manufacturer’s instructions.

### Culture and differentiation of THP1 monocytes

THP1s were cultured in RPMI (Gibco) supplemented with 10% FBS, GlutaMax, and 1% penicillin and streptomycin. 1.5 x 10^6^ cells were plated in a 12-well plate with 50 ng/mL of phorbol 12-myristate-13-acetate (PMA; Sigma). After 24 hours, media and non-adherent cells were removed and replaced with fresh RPMI prepared as described above. Cells were rested for an additional 24 hours before treatment.

### Murine bone marrow derived macrophage culture

Femur and tibia bones were removed from 8-12-week-old mice, and bone marrow was flushed using fresh RPMI media. Bone marrow was cultured in RPMI-1640 media (ATCC) supplemented with 10% FBS, 1% penicillin and streptomycin, and 20% L929 conditioned media produced as described above. Cells were incubated at 37 °C in a humidified atmosphere of 5% CO_2_ for 7 days.

### Cas9 gene knockdown in murine BMDMs

BMDM were isolated and differentiated as described above. Pooled guides (EditCo) were reconstituted per manufacturer’s instructions. P3 Primary Cell Nucleofector Solution and Supplement 1 (Lonza) were combined per manufacturer’s instructions immediately before use. The 20 pmol Cas9 (EditCo) and 180 pmol sgRNA was precomplexed at room temperature for 20 mins – 1 hour in P3 solution to generate RNP. BMDMs were lifted, washed with PBS, and resuspended in P3 solution, nucleofecting 2.5 x 10^6^ cells per reaction. Mix the RNP and cells by gently pipetting and load into Nucleocuvette Strip (Lonza). Strips were immediately inserted into the Lonza 4D-nucleofector (Lonza) and nucleofected with the Buffer P3, CM-137 condition. Cells were immediately resuspended in warm complete RPMI media containing 5% LCCM and plated at a concentration of 1.5 x 10^6^ cells/mL. The media was exchanged 24 hours after nucleofection, and cells were allowed to rest for 72 hours before treatments and assays were performed. Knockout efficiency was assessed via immunoblotting.

### Isolation and culture of human monocyte-derived macrophages

PBMCs were isolated using Lymphoprep (StemCell) and stored in liquid nitrogen until use. Monocytes were isolated using a CD14-positive selection kit (Miltenyi) and LS positive selection columns (Miltenyi) per manufacturer’s instructions. After isolation, monocytes were resuspended at 1 x 10^6^ cells/mL in complete RPMI-1640 media (ATCC RPMI, 10% FBS, 1 U/mL penicillin, 100 mg/mL streptomycin) supplemented with 50 ng/mL M-CSF (Biolegend). Cells were plated at 2 x 10^6^ cells/well in 6-well plate (Sigma) and incubated at 37°C. 1 day after isolation, each well was supplemented with an additional 2 mL of complete RPMI media containing M-CSF, and 4 days after isolation, 50% of the media was exchanged. 7 days after isolation, cells were washed with PBS and lifted from the plate with 2 mM EDTA in PBS, counted, and plated in M-CSF - supplemented media at 1 x 10^6^ cells per well in a 12-well tissue culture-treated plate, or 1 x 10^5^ cells/well in a 96 well plate.

### Isolation of peritoneal macrophages

Mice were injected IP with 3 mL of sterile, aged (3-4 weeks) 3% thioglycolate (Sigma). 4 days after administration of thioglycolate, peritoneal macrophages were isolated via flushing the peritoneal cavity twice with 10 mL of cold, sterile PBS. To isolate cells, the peritoneal lavage fluid was centrifuged (300 x g for 5 mins), and red blood cells were removed with RBC lysis buffer (Sigma). 5 x 10^5^ cells were plated on a poly-L-lysine coated glass coverslip (EMS) and rested at 37°C overnight before being fixed and stained as described.

### Flow cytometry

Flow cytometry was performed as previously described (*48*). Briefly, BMDMs were stained in MACS buffer (0.5% BSA, 2 mM EDTA) using Ghost Violet 540 viability dye (TONBO), anti-CD45.2 BV650 (BioLegend), CD11b PE (BioLegend), and MARCO APC (eBioscience). Cells were acquired on a CytekTM Aurora. Flow cytometry analysis was done using FlowJo software.

### RNA sequencing

RNA sequencing was performed by BGI as previously described (*49*).

### Peptide mapping by nano LC-MS/MS

Streptavidin pull downs were eluted in pierce elution buffer and subjected to in solution to in-gel digestion with trypsin. The resulting peptides were lyophilized, re-suspended in 5% acetonitrile, 0.1% (v/v) formic acid in water and injected onto a NanoAcquity UPLC (Waters) coupled to a Q Exactive (Thermo Scientific) hybrid quadrapole orbitrap mass spectrometer. Peptides were trapped on a 100 µm I.D. fused silica pre-column packed with 2 cm of 5 µm (200Å) Magic C18AQ (Bruker-Michrom) particles in 5% acetonitrile, 0.1% (v/v) formic acid in water at 4.0 µl/min for 4.0 minutes. Peptides were then separated over a 75 µm I.D. gravity-pulled 25 cm long analytical column packed with 3 µm (100Å) Magic C18AQ particles, at a flow rate of 300 nl/min containing mobile phase A, 0.1% (v/v) formic acid in water and mobile phase B, 0.1% (v/v) formic acid in acetonitrile, using a biphasic gradient: 0-60 min (5-35% B), 60-90 min (35-60% B), 90-93 min (60% B), 93-94 min (60-90% B), 94-109 (90% B), followed by equilibration to 5% B. Nano-ESI source was operated at 1.4 kV via liquid junction. Mass spectra were acquired over m/z 300-1750 at 70,000 resolution (m/z 200) with an AGC target of 1e6. Data-dependent acquisition (DDA) selected the top 10 most abundant precursor ions for tandem mass spectrometry by HCD fragmentation using an isolation width of 1.6 Da, max fill time of 110ms, and AGC target of 1e5. Peptides were fragmented by a normalized collisional energy of 27, and product ion spectra were acquired at a resolution of 17,500 (m/z 200). Raw data files were peak processed with Proteome Discoverer (version 2.1, Thermo Scientific) followed by identification using Mascot Server (Matrix Science) against the Mouse (Swissprot) FASTA file (downloaded 07/2019). Search parameters included full tryptic enzyme specificity, and variable modifications of N-terminal protein acetylation, oxidized methionine, glutamine conversion to glutamic acid. Assignments were made using a 10-ppm mass tolerance for the precursor and 0.05 Da mass tolerance for the fragment ions. All non-filtered search results were processed by Scaffold (version 4.8.4, Proteome Software, Portland, OR) utilizing the Trans-Proteomic Pipeline (Institute for Systems Biology, Seattle, WA) at 1% false-discovery rate (FDR) for peptides and 99% threshold for proteins (2 peptides minimum).

### Immunofluorescence

Coverslips were handled with sterile forceps, retrieved from -20°C storage, and placed in 24-well plates. BMDM cells were seeded at 5 x 10^5^ cells per poly-L-Lysine glass cover slip (EMS) placed in a 24 well plate incubated overnight before treatments. Following treatment, media was removed, and cells were washed with PBS. Cells were fixed in 4% paraformaldehyde (Fisher) for and washed in PBS. Cells were permeabilized with 0.1% Tween-20. Cells were blocked with PBS supplemented with 5% normal goat serum (Sigma) and 0.3% triton (Sigma). Coverslips were incubated in primary antibodies (MARCO, 1:100 (Abcam), NRF2, 1:100 (Cell Signaling), p62, 1:200 (Cell Signaling) diluted in blocking buffer. Coverslips were incubated in secondary antibody (anti-rabbit, 1:200 (ThermoFisher)) in PBS. Cells were incubated in 1 μg/mL Hoechst (Thermo Scientific). Coverslips were carefully removed and placed on slides prepared with a drop of DAPI-containing mounting solution (ThermoFisher) onto microscope slides (Fisher) and cured overnight before imaging. Slides were imaged with the Leica Thunder microscope quantification was performed using ImageJ/Fiji.

### LPS Septic Shock

Mice were intraperitoneally injected with 40mg/kg of LPS (Sigma, stock concentration 5 mg/mL) or equivalent volume of vehicle (water). Survival was then monitored over time.

### Polymicrobial Sepsis

Cecal slurry stock was isolated as previously described (*50*). Briefly, whole cecal content dissection from WT mice bred in house were weighed and resuspended in 0.5 mL 1x PBS per 100 mg of cecal content. Slurry was vortexed until homogeneous and filtered through a 100 μm filter. Equal volumes of 30% glycerol/PBS were added, and glycerol stocks were aliquoted and frozen down in −80°C until use. Cecal Slurry shock was conducted by quick thawing cecal slurry aliquot at 37°C and 150 μL of slurry was injected via intraperitoneal injection.

### *Salmonell*a Typhimurium Infection

Cultures of *Salmonella* Typhimurium strain SL1344 were grown in LB broth (Sigma) supplemented with 100 μg/mL streptomycin (Sigma) for 16 hours overnight at 37°C, shaking at 200 rpm. The overnight culture was diluted into streptomycin-containing LB and incubated at 37°C, shaking at 200 rpm for 2 – 4 hours, until OD_600_ = 1.0 (approximately 1 x 10^9^ CFU/mL). Culture was serially diluted from 1 x 10^-5^ to 1 x 10^-9^ and plated on LB agar plates (Sigma) containing 100 μg/mL streptomycin. Plates were incubated overnight at 37°C to determine the inoculation CFU as determined by the formula: # colonies x dilution factor / plating volume (mL) = CFU/mL. For the inoculation, approximately 1 x 10^3^ CFU per mouse was isolated, washed 2x in PBS, and resuspended at 1 x 10^5^ CFU/mL in PBS. 100 μL of the inoculum was immediately injected IP. Mice were monitored for weight loss and survival.

### Isolation of peritoneal macrophages for assessing bacterial burden

Peritoneal macrophages were isolated via peritoneal lavage as previously described. Following isolation of the peritoneal cells, the lavage fluid supernatant was collected for assessing bacterial burden. 5 x 10^5^ cells were plated on a poly-L-lysine coated glass coverslip (EMS) and rested at 37 °C overnight, and fixed and stained as described. The peritoneal lavage fluid was serially diluted up to 1 x 10^-3^ and plated on LB agar plates containing 100 μg/mL streptomycin. Plates were incubated overnight at 37 °C and CFU/mL was determined as described above.

### Genotyping

Ear clips collected from mice were incubated with 50 nM NaOH for 1 hour at 55 °C to dissociate tissue. The DNA digest was then used for PCR, using the following primer sequences, and the amplicon length was determined by via electrophoresis.

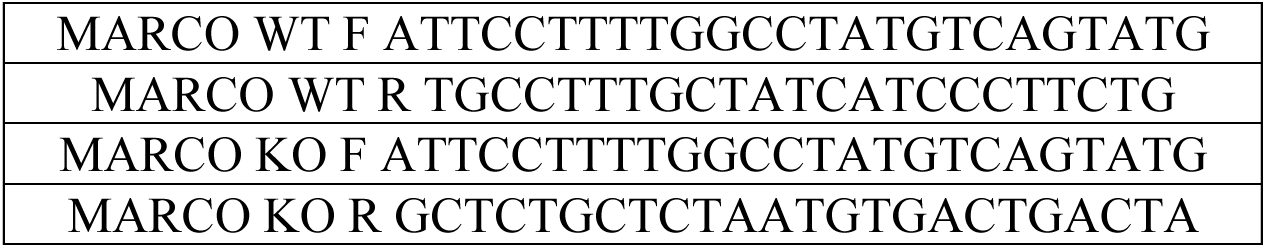

### Histology (spleen tissue)

Processing and mounting of mouse spleens was performed by Applied Pathology Systems, LLC. Tissues were fixed in 10% formalin (Sigma) then transferred to 70% ethanol. The fixed tissues were dehydrated and cleared with xylene to remove alcohol. Tissues were subjected to paraffin infiltration and embedding for sectioning. Thin tissue sections, 4–5 µm, were sectioned with microtome and mounted on glass slides (Fisher) and dried in an oven at 60°C for 1 hour. Antigen retrieval was performed on dewaxed and rehydrated FFPE sections using R-Buffer A (Fisher Scientific) using 2100 antigen retriever (Aptum Biologics Ltd). For immunofluorescence staining, tissue sections were blocked for 1 hour with 5% donkey serum (Sigma) in 0.3% Triton-X (Sigma) in PBS. The tissue sections were incubated with primary antibodies (MARCO, 1:200 (Abcam), F4/80 1:200 (ThermoFisher) diluted in blocking buffer. Following incubations, slides were washed in PBS containing 0.1% Tween (Sigma). Tissue sections were incubated with secondary antibodies (anti-rabbit, 1:200 (ThermoFisher), anti-rat, 1:200 (ThermoFisher)) diluted in PBS. The slides were mounted using DAPI containing Fluro mount-G Mounting media (ThermoFisher) with cover glasses (Sigma). Slides were imaged with the Leica Thunder microscope.

### Statistical analysis

For comparisons of two groups two-tailed students’ t test was performed. Multiple comparison analysis was performed using two-way ANOVA. Mantel-Cox was used for survival analysis. Three to ten mice were used per experiment, sufficient to calculate statistical significance, and in line with similar studies published in the literature.

### Ethics

All animal studies were performed in compliance with the federal regulations set forth in the Animal Welfare Act (AWA), the recommendations in the Guide for the Care and Use of Laboratory Animals of the National Institutes of Health, and the guidelines of the UMass Medical School Institutional Animal Use and Care Committee. All protocols used in this study were approved by the Institutional Animal Care and Use Committee at the UMass Medical School (protocol 202200082).

## Acknowledgements

FH is funded by start-up funding from UMass Chan Medical School, Charles H.Hood foundation, Riccio Fund for neuroscience, the UMass Chan BRIDGE fund, and NIAID R01AI199543. SC was funded by NIH/NIAID T32 AI095213 and F31 AI197725. We thank Prof. Katherine Fitzgerald for *Ifnar*-/- and *Irf3*^-/-^ mice.

## Author Contributions

SR and SC performed experiments, analyzed data, and edited the manuscript. MF assisted with animal experiments and animal husbandry. AS and KC isolated and cultured primary human macrophages. LS assisted with experiments. LG performed experiments and analyzed data and edited the manuscript. FH conceived the study, developed the concept, performed experiments, analyzed data and wrote the manuscript.

## Supplementary Information

**Supplementary Figure 1:**
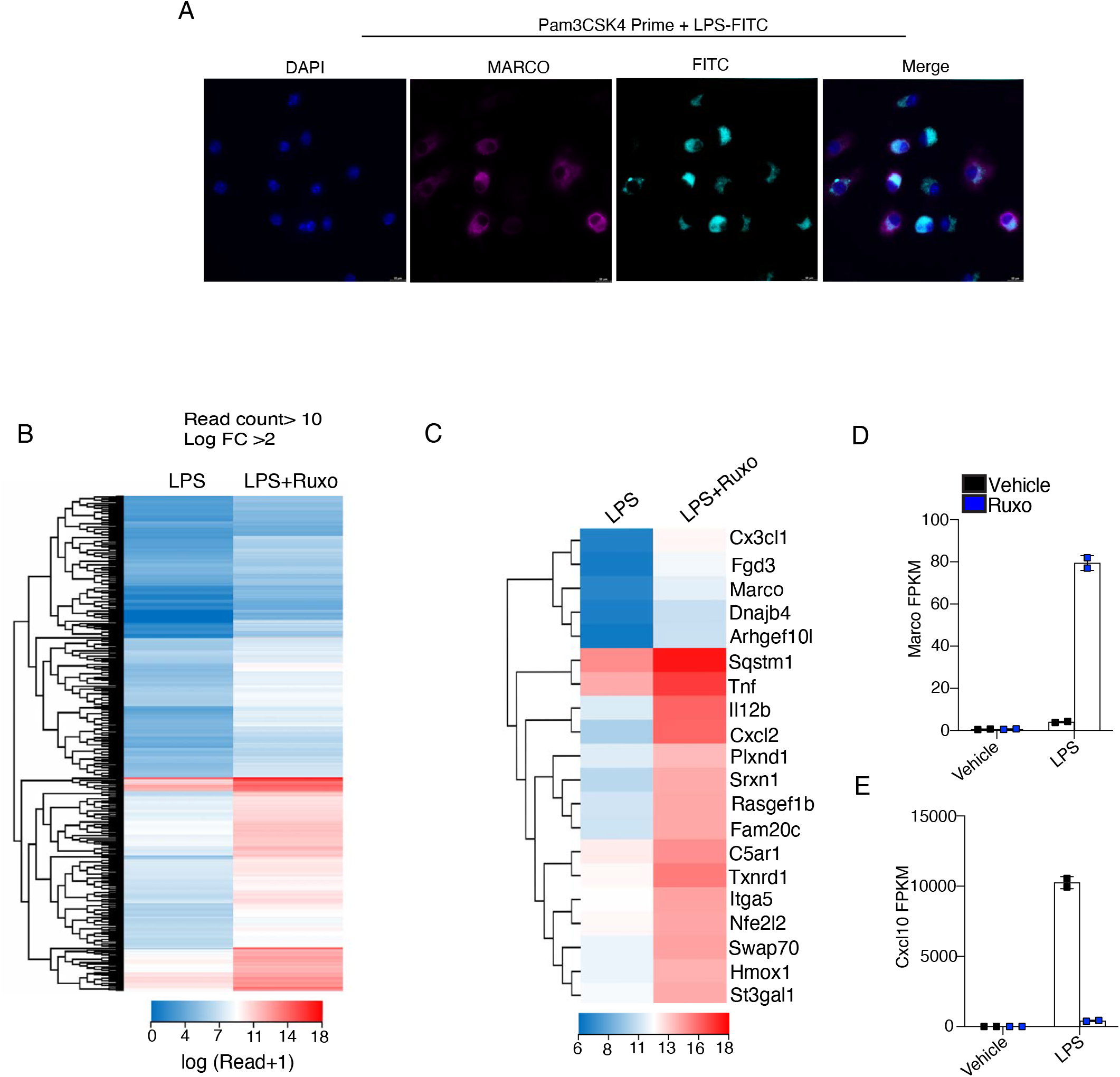
RNA sequencing analysis of ruxolitinib treated macrophages. **(A)** Immunofluorescence staining of MARCO (pink), FITC-labelled LPS (cyan), and Hoechst (blue) in WT BMDMs primed with Pam3CSK4 (1μg/mL) for 24 hours followed by incubation with FITC-LPS (500ng/mL) for 1 hour. **(B)** Heat map of differentially expressed genes from RNA-seq analysis of vehicle or ruxolitinib (1μM) and LPS (100ng/mL) treated BMDMs. **(C)** Top 20 most significantly upregulated genes in LPS (100ng/mL) and ruxolitinib (1μM) treated cells relative to LPS (100ng/mL) treated cells. (**D-E)** FPKM values of *Marco* (D) and *Cxcl10* (E). A representative image of 3 independent experiments. B-D average of two-independent experiments from RNA sequencing.

**Supplementary Figure 2:**
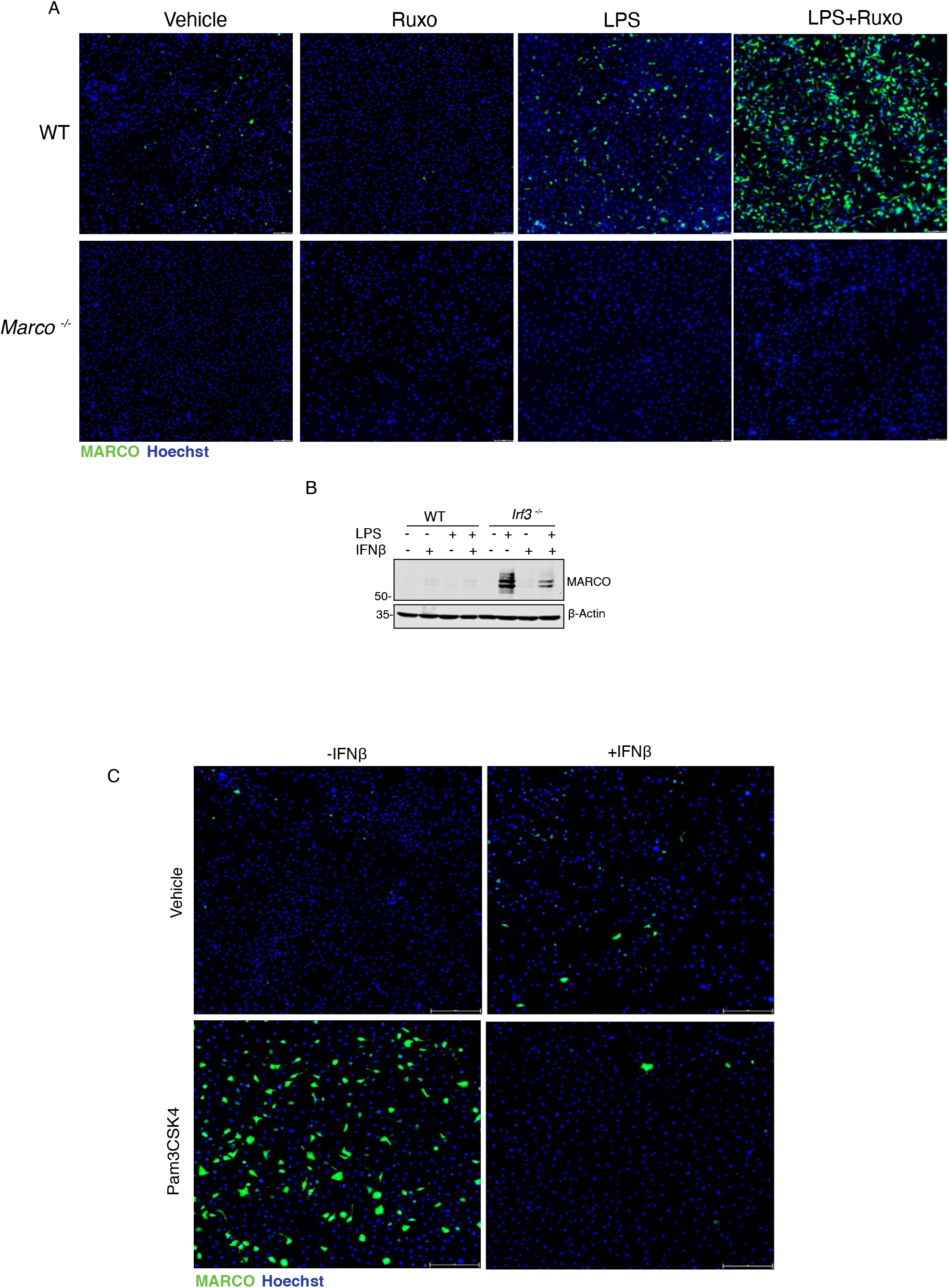
IFN negatively regulates MARCO expression. **(A)** Immunofluorescence staining of MARCO (green) and Hoechst (blue) in WT and *Marco ^-/-^* BMDMs treated with by ruxolitinib (1μM) for 1 hour followed by LPS (100ng/mL) for 24 hours. **(B)** Immunoblot of MARCO and β-actin in WT and *Irf3 ^-/-^* BMDMs treated LPS (100ng/mL) for 1 hour followed by IFNβ (10ng/mL) for 24 hours. **(C)** Immunofluorescence staining of MARCO (green) and Hoechst (blue) in WT BMDMs treated with Pam3CSK4 (1μg/mL) for 1 hour followed by IFNβ (50ng/ml) for 24 hours. A-C representative images from 3 independent experiments.

**Supplementary Figure 3:**
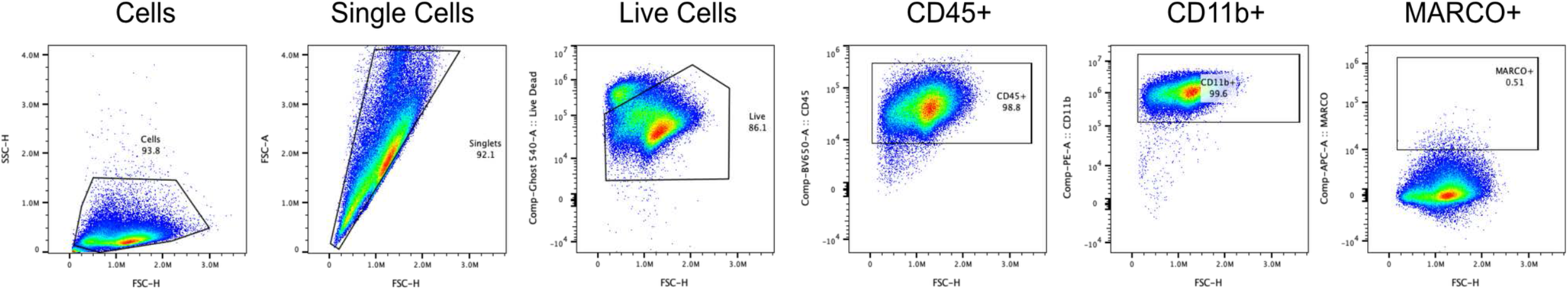
Flow cytometry gating strategy for Figure 3. Representative flow plots for the gating strategy of single live CD45+ CD11b+ MARCO+ cells. Representative plots from 3 independent experiments.

**Supplementary Figure 4:**
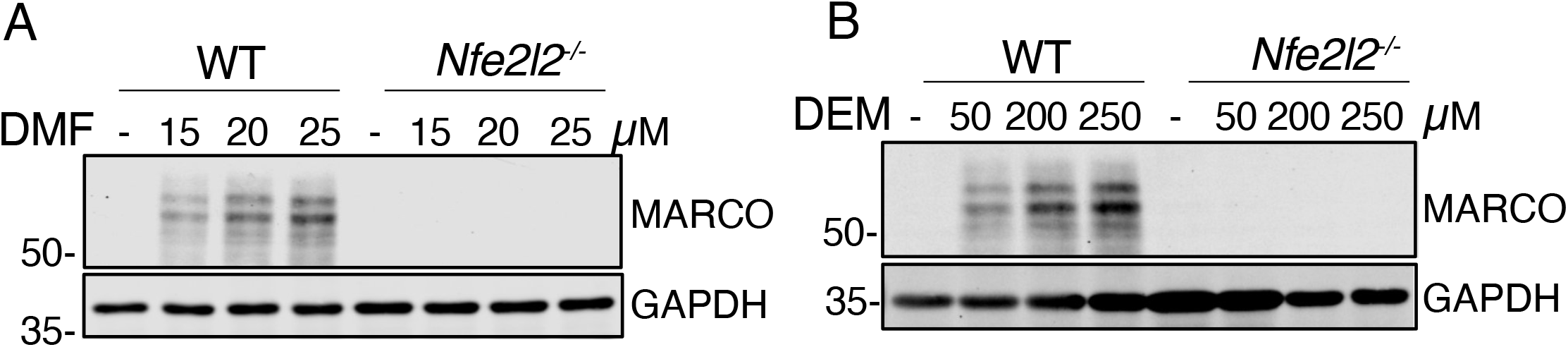
NRF2 activators induce MARCO expression. **(A-B)** Immunoblot for MARCO and GAPDH in WT and *Nfe2l2^-/-^*BMDMs treated with DMF (A) or DEM (B) at the indicated concentrations for 24 hours. A-B representative images from 3 independent experiments.

**Supplementary Figure 5:**
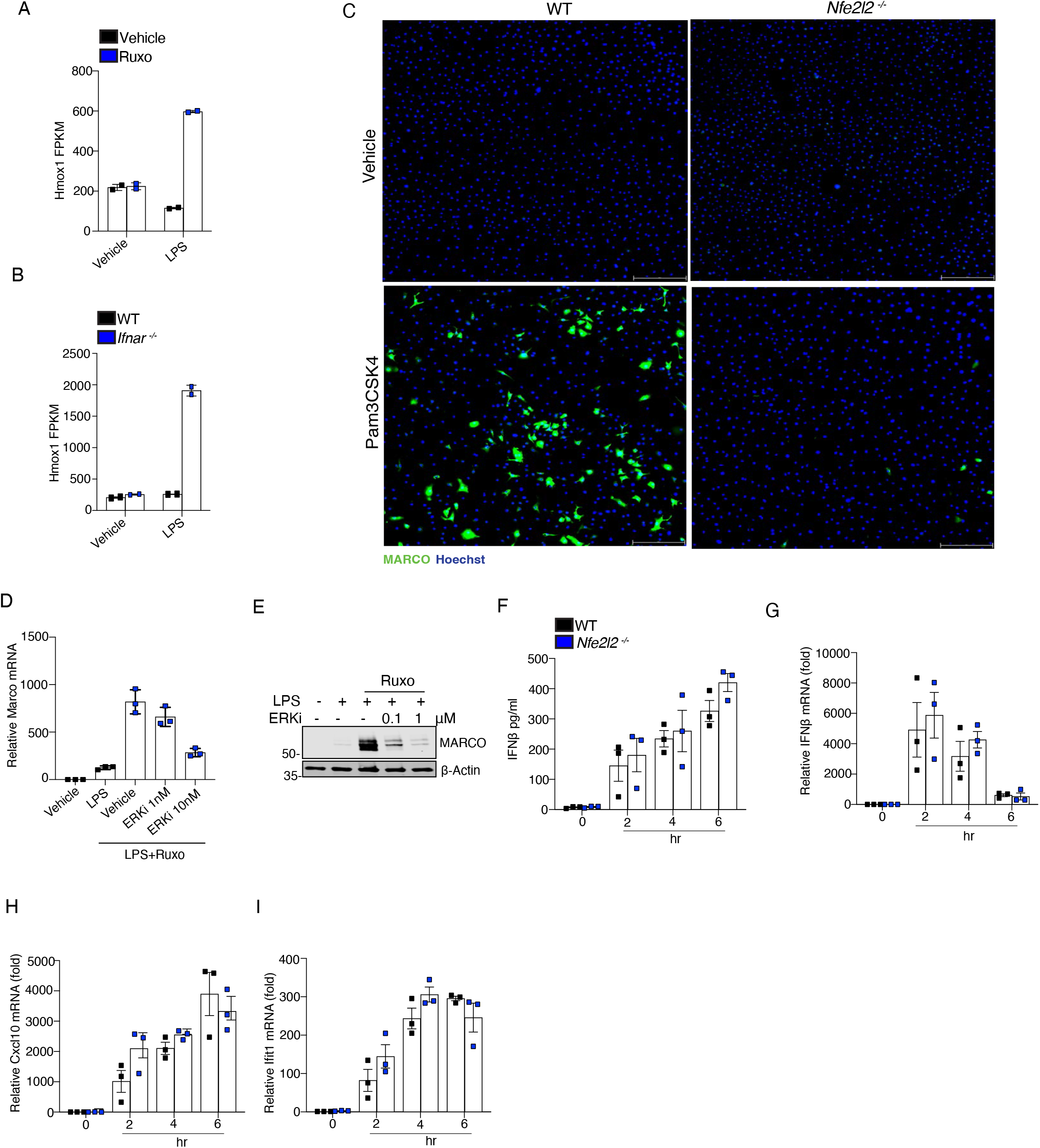
IFN negatively regulates NRF2 dependent gene expression. **(A-B)** FPKM values of *Hmox1* in LPS (100 ng/mL) and ruxolitinib (1μM) treated cells (A) and WT and *Ifnar ^-/-^* BMDMs treated with LPS (100ng/mL) for 6 hours (B). **(C)** Immunofluorescence staining of MARCO (green) and Hoechst (blue) in WT and *Nfe2l2 ^-/-^* BMDMs treated Pam3CSK4 (1μg/mL) for 24 hours. **(D)** RT-qPCR analysis of *Marco* mRNA in WT BMDMs treated with or without AZD0364 (ERKi) at the indicated concentrations for 1 hour followed by treatment with LPS (100ng/mL) and ruxolitinib (1μM) for 6 hours. **(E)** Immunoblot of MARCO and β-actin in lysates from WT BMDMs treated with or without AZ0364 (ERKi) at the indicated concentrations for 1 hour, followed by treatment with LPS (100ng/mL) and ruxolitinib (1μM) for 24 hours. **(F)** ELISA analysis of IFNβ in the supernatant of WT and *Nfe2l2^-/-^* BMDMs following treatment with LPS (100ng/mL) for the indicated time points. **(G-I)** RT-qPCR analysis of *Ifnβ* (G), *Cxcl10* (H), and *Ifit1*(I) mRNA transcripts following treatment of WT and *Nfe2l2^-/-^* BMDMs with LPS (100ng/mL) for the indicated time points. A-B average of two-independent experiments from RNA sequencing described in Figure 2 and Extended Figure 1. C, E representative images from 3 independent experiments. D, F-I, pooled biological replicates from 3 independent experiments.

**Supplementary Figure 6:**
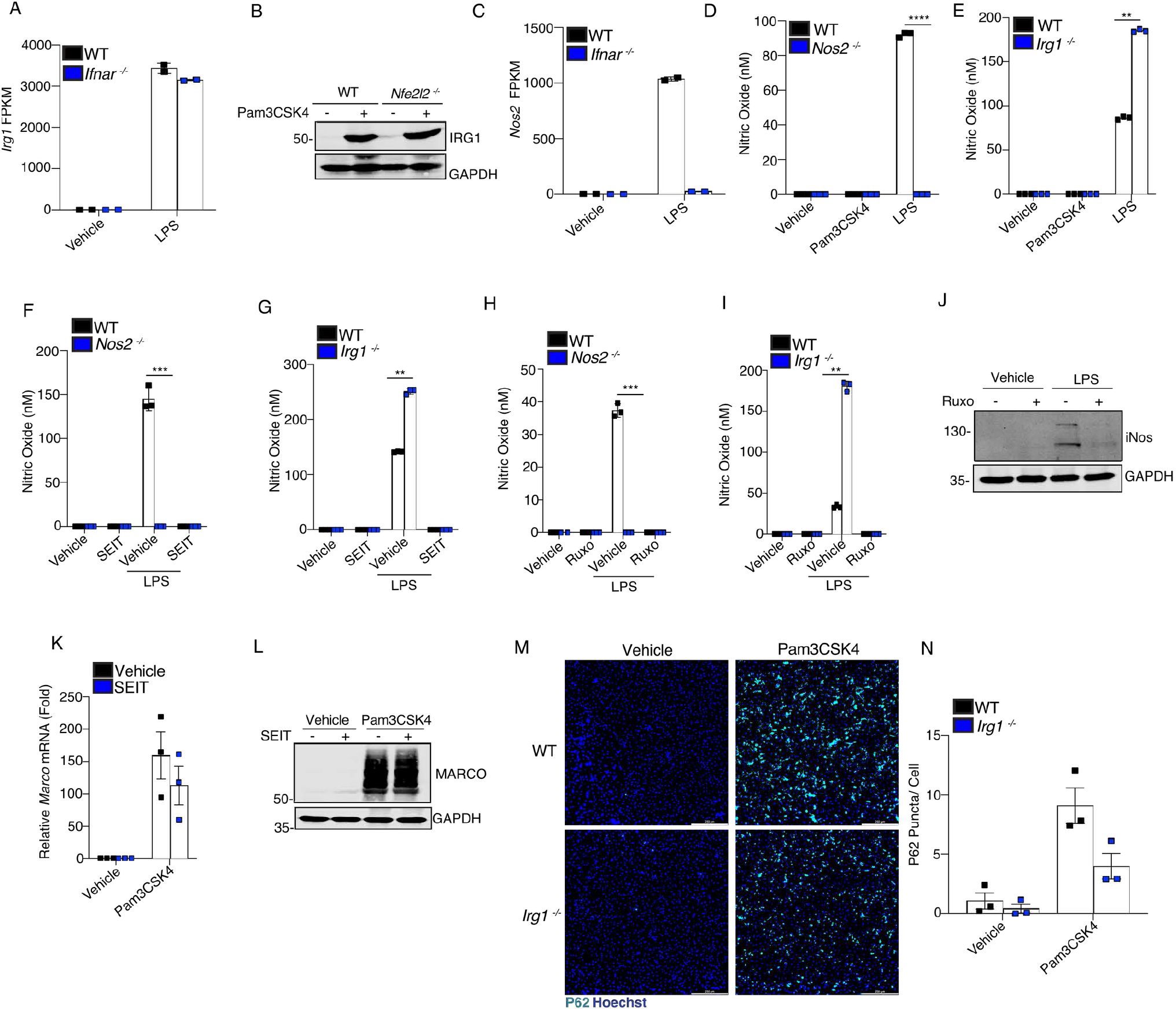
Nitric oxide is not required for MARCO expression. **(A)** FPKM values of *Irg1* in RNA seq of WT and *Ifnar* ^-/-^ BMDMs treated with vehicle or LPS (100ng/mL) for 6 hours. **(B)** Immunoblot analysis of IRG1 and GAPDH in WT and *Nfe2l2* ^-/-^ BMDMs with Pam3CSK4 (1μg/ml) for 24 hours. **(C)** FPKM values of *Nos2* from RNA sequencing analysis of WT and *Ifnar ^-/-^* BMDMs treated with vehicle or LPS (100ng/mL) for 6 hours. **(D-E)** Nitric oxide assay on supernatants from WT and *Nos2* ^-/-^ (D) or WT and *Irg1 ^-/-^* (E) treated with Pam3CSK4 (1μg/mL) or LPS (100ng/mL) for 24 hours. **(F-G)** Nitric oxide assay on supernatants from WT and *Nos2 ^-/-^* (F) or WT and *Irg1 ^-/-^* (G) BMDMs treated with or without LPS (100ng/mL) with or without SEIT (250mM) for 24 hours. **(H-I)** Nitric oxide assay on supernatants from WT and *Nos2* ^-/-^ (H) or WT and *Irg1 ^-/-^*(I) treated with or without LPS (100ng/mL) with or without ruxolitinib (1μM) for 24 hours. **(J)** Immunoblot of iNOS and GAPDH in lysates from WT BMDMs treated with or without ruxolitinib (1μM) for 1 hour, followed by treatment with LPS (100ng/mL) for 24 hours. **(K)** RT-qPCR analysis of *Marco* mRNA in WT BMDMs treated with or without SEIT (250nM) for 1 hour, followed by Pam3CSK4 (1μg/mL) for 5 hours. **(L)** Immunoblot of MARCO and GAPDH in lysates from WT BMDMs treated with or without SEIT (250mM) for 1 hour, followed by treatment with Pam3CSK4 (1μΜ) for 24 hours. **(M-N)** Immunofluorescence (M) and quantification (N) of p62 (cyan) aggregation and Hoechst (blue) in WT and *Irg1^-/-^*BMDMs treated with Pam3CSK4 (1μg/mL) for 24 hours. A, C average of two-independent experiments from RNA sequencing described in Figure 2. B, J, L, M, representative images from 3 independent experiments. D-I, K, N pooled biological replicates from 3 independent experiments.

**Supplementary Figure 7:**
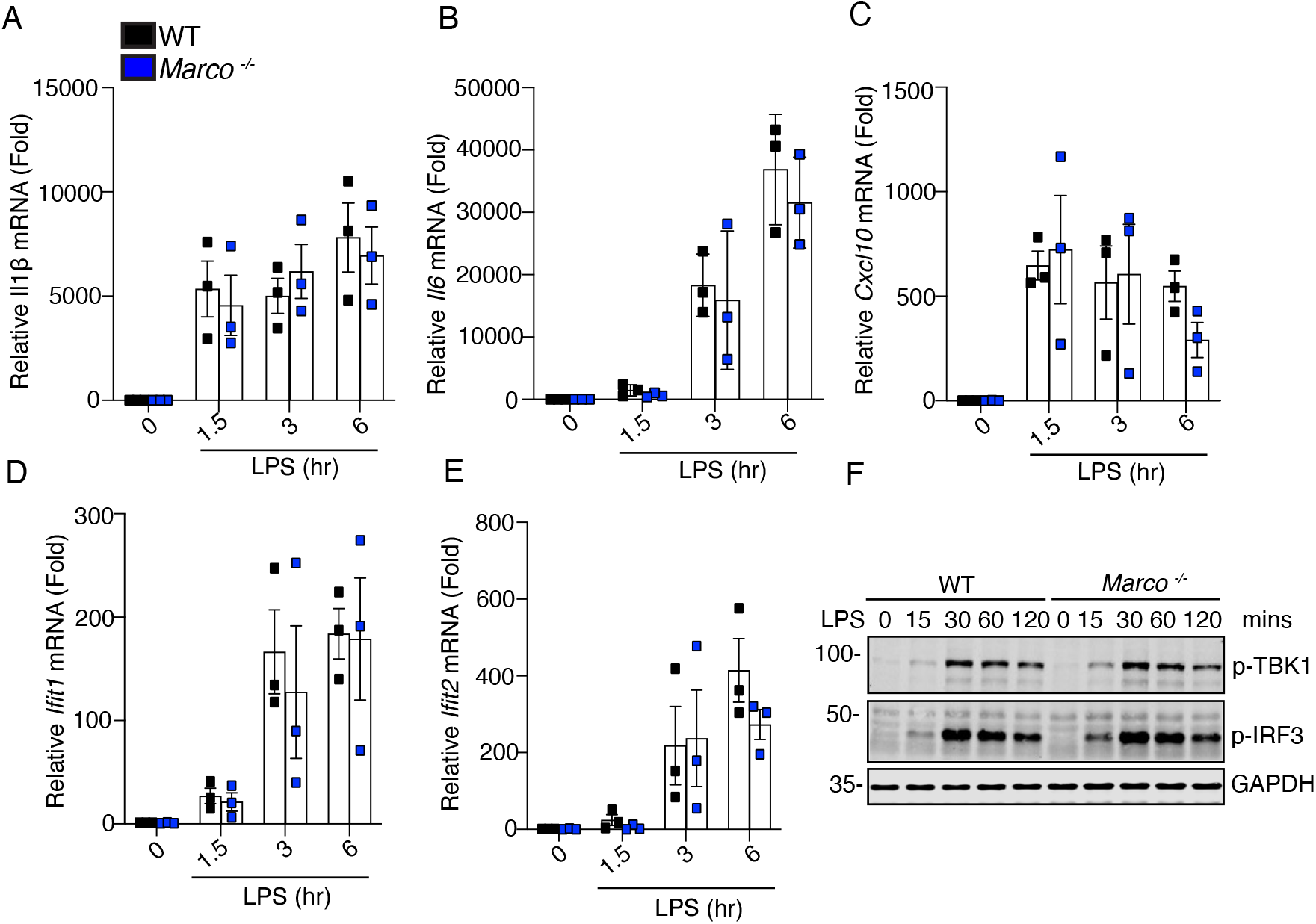
MARCO is dispensable for TLR4 signaling in BMDMs. **(A-E)** RT-qPCR analysis of *Il1β* (A)*, Il6* (B)*, Cxl10* (C)*, Ifit1* (D)*, Ifit2* (E) mRNA in WT and *Marco* ^-/-^ BMDMs treated with LPS (100ng/mL) for the indicated times. **(F)** Immunoblot analysis of phosphorylated IRF3 and TBK1 and GAPDH in lysates from WT and *Marco* ^-/-^ BMDMs treated with LPS (100ng/mL) for the indicated times. A-E pooled biological replicates from 3 independent experiments. F, representative images from 3 independent experiments.

**Supplementary Figure 8:**
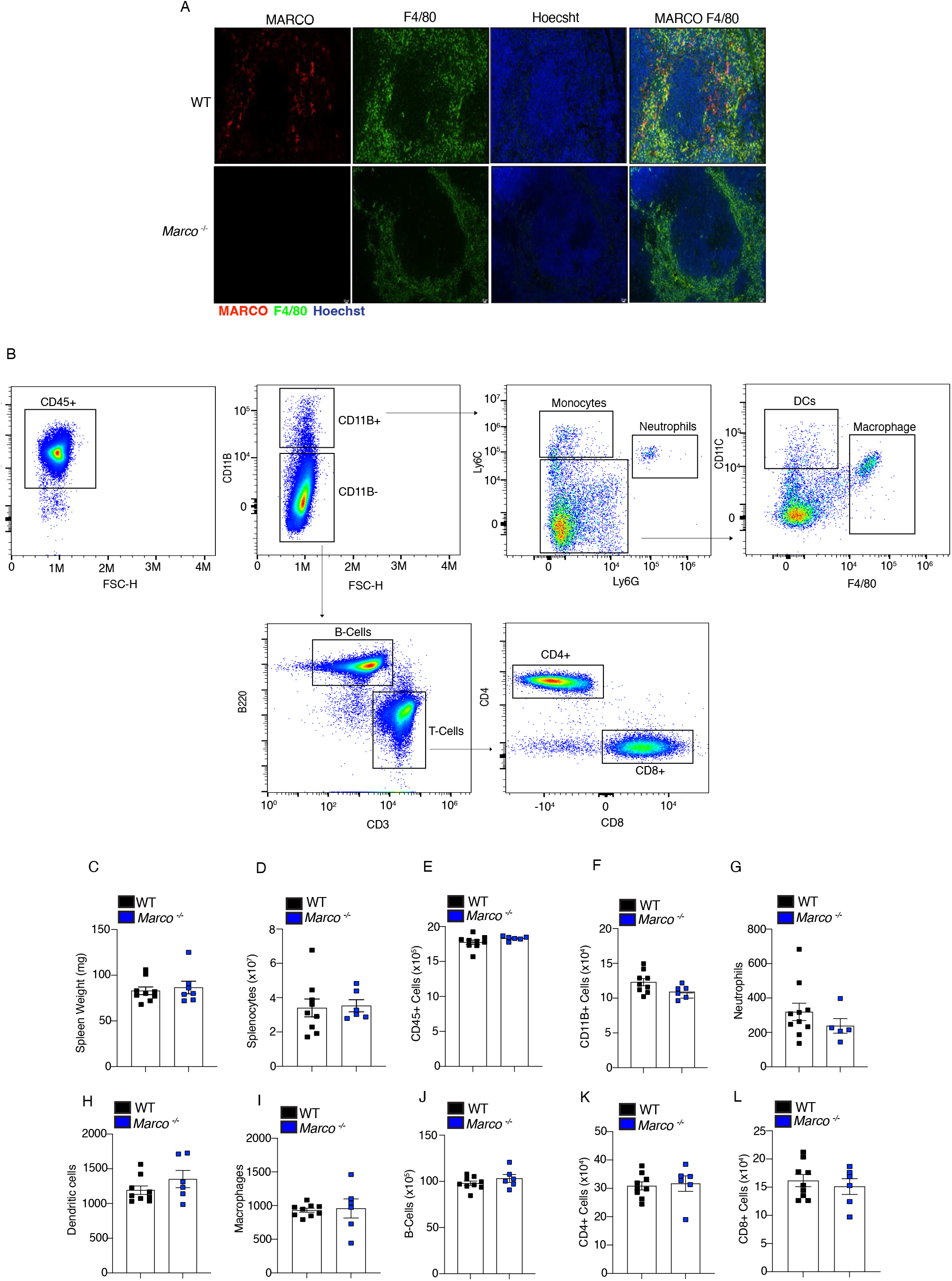
Splenic immune cell profile of *Marco*-deficient mice. **(A)** Immunofluorescence staining of MARCO (red), F4/80 (green), and Hoechst (blue) from splenic tissue sections from WT and *Marco ^-/-^* mice. **(B)** Representative flow plots and gating strategy of live, single, CD45+ cells. **(C-D)** Spleen weight (C) and splenocyte number (D) of spleens from WT and *Marco* ^-/-^ mice. **(E-L)** Cell counts of the indicated cell types in spleens from WT and *Marco* ^-/-^ mice. A representative image of 3 independent experiments. B is representative plots of n=5-9 mice, C-L is pooled data from *n*=5-9 mice per group.

**Supplementary Figure 9:**
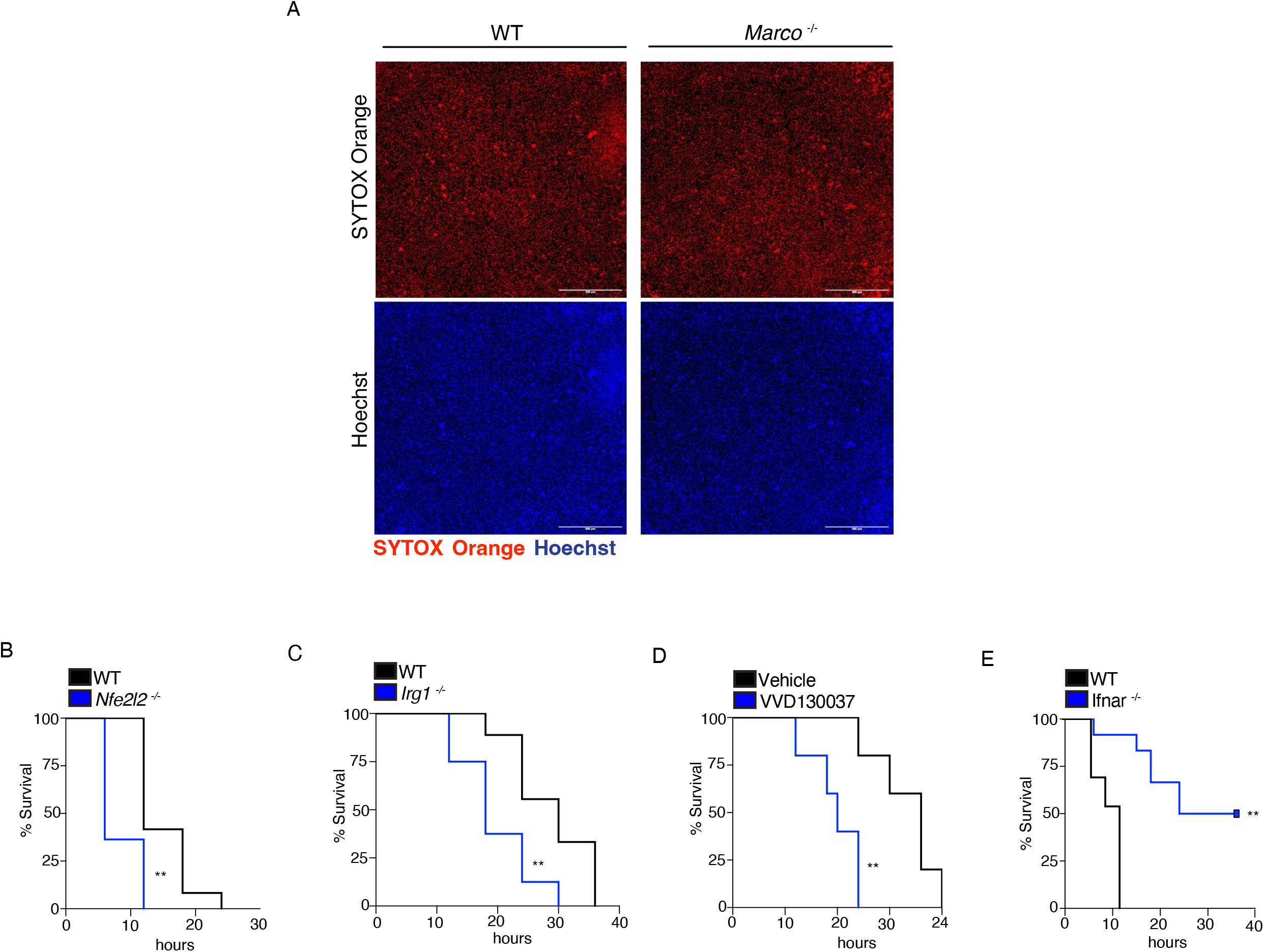
Loss of NRF2 or itaconate sensitizes mice to LPS shock. **(A)** Representative images of SYTOX+ (orange) and Hoechst (blue) cells of WT and *Marco ^-/-^* BMDMs treated with LPS (100ng/mL) for 3 hours and nigericin (10μM) for 1 hour. **(B)** Survival analysis of WT and *Nfe2l2* ^-/-^ mice administered LPS (40mg/kg). **(C)** Survival analysis of WT and *Irg1*^-/-^ mice administered LPS (40mg/kg). **(D)** Survival analysis of WT mice administered LPS (40mg/kg) with or without pre-treatment with VVD130037 (10mg/kg). **(E)** Survival analysis of WT and *Ifnar* ^-/-^ mice administered LPS (40mg/kg). A representative image of 3 independent experiments. B-E is pooled from n=5-10 mice. **P<0.01, Mantel-Cox analysis

**Supplementary Figure 10:**
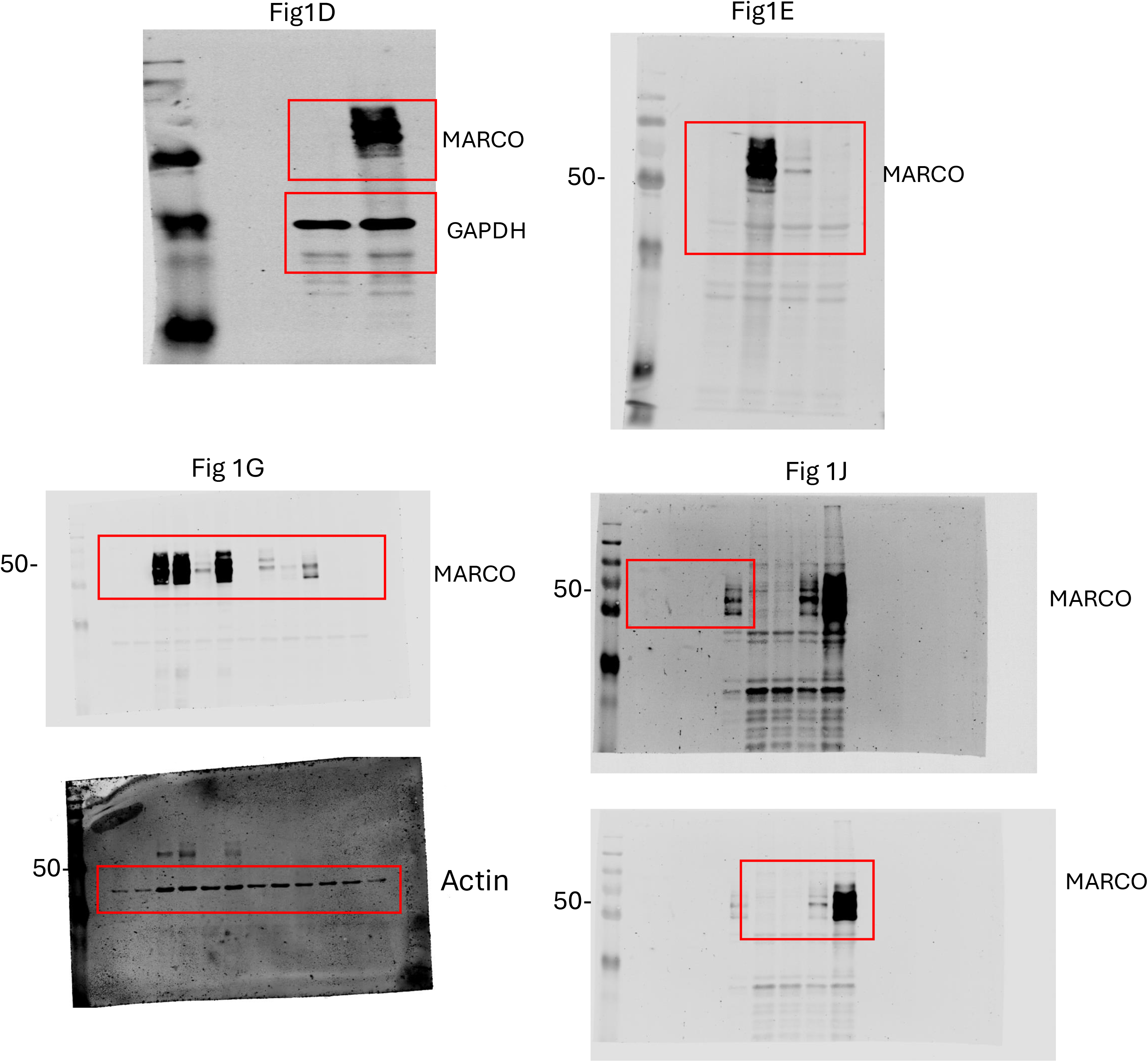
Uncropped western blots from Figure 1.

**Supplementary Figure 11:**
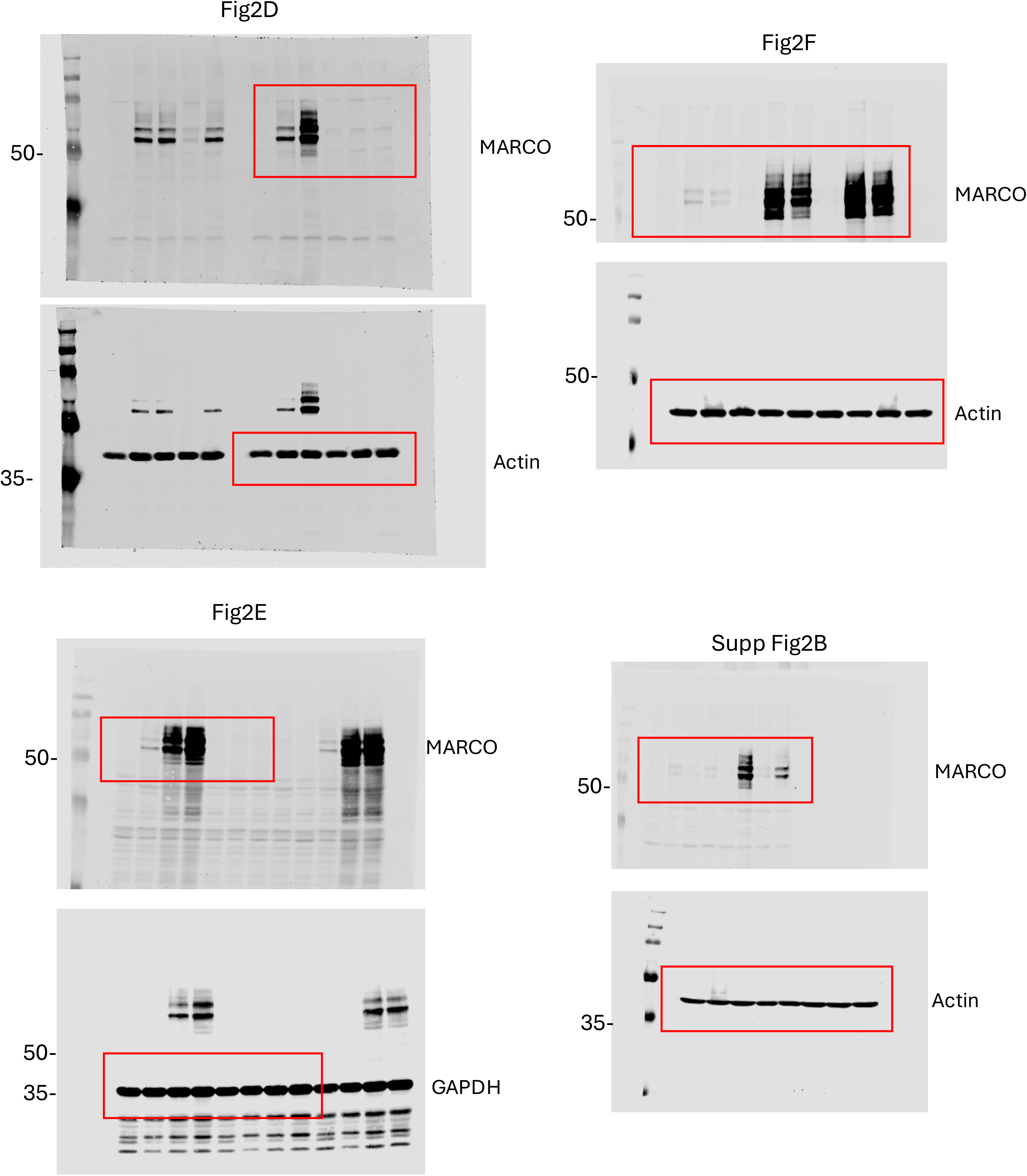
Uncropped western blots from Figure 2 and Supp Figure 2.

**Supplementary Figure 12:**
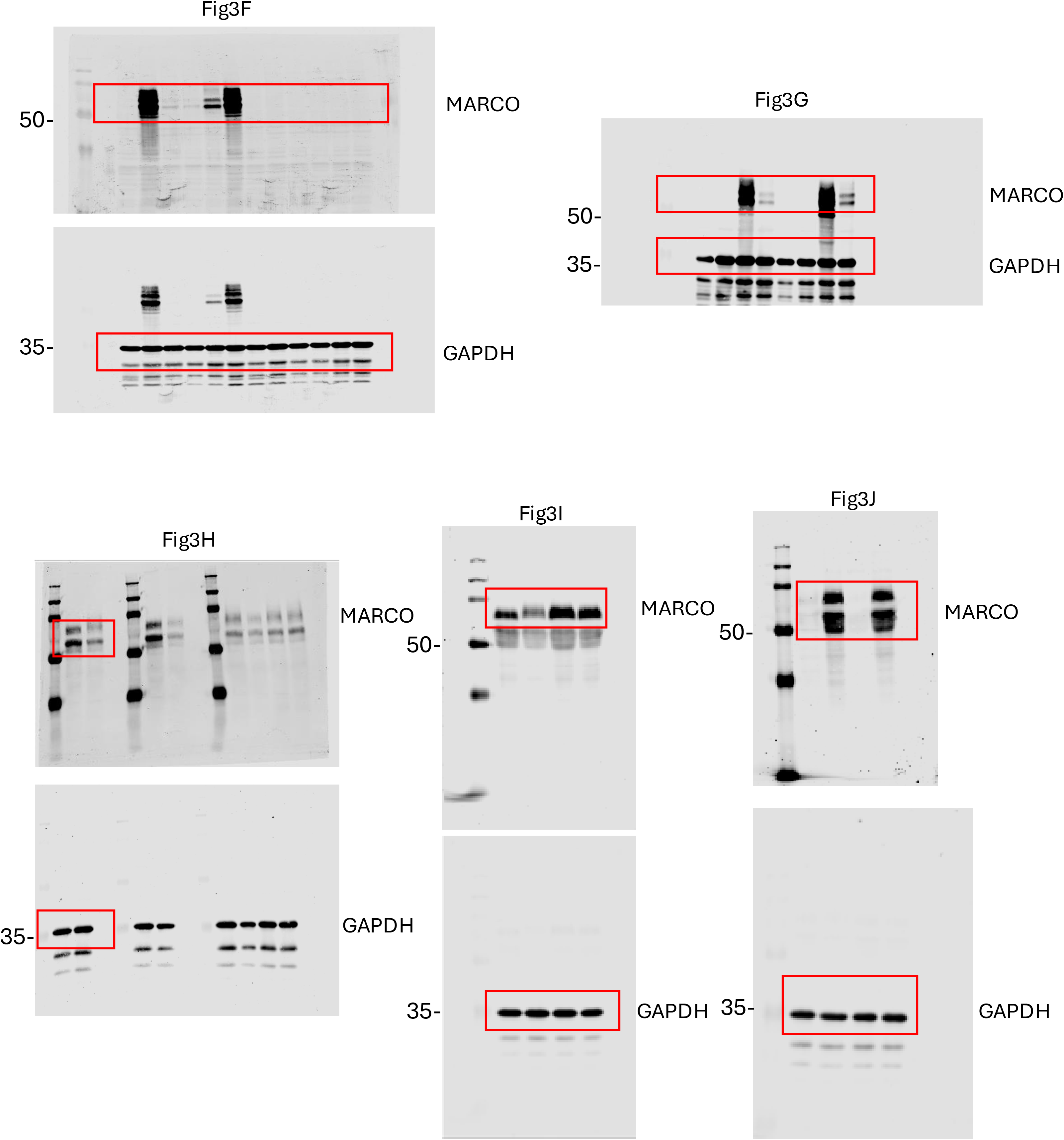
Uncropped western blots from Figure 3.

**Supplementary Figure 13:**
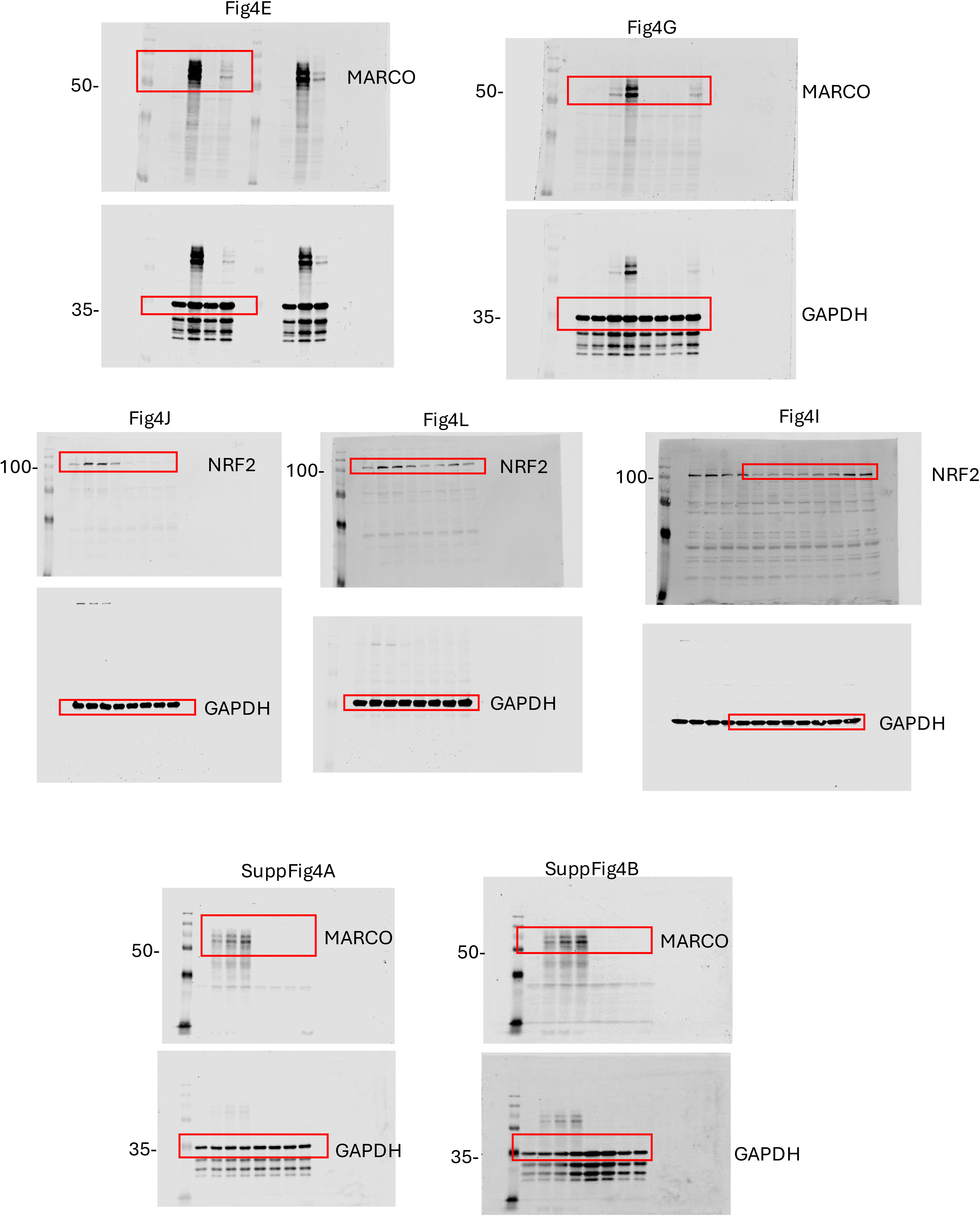
Uncropped western blots from Figure 4 and supp Figure 4.

**Supplementary Figure 14:**
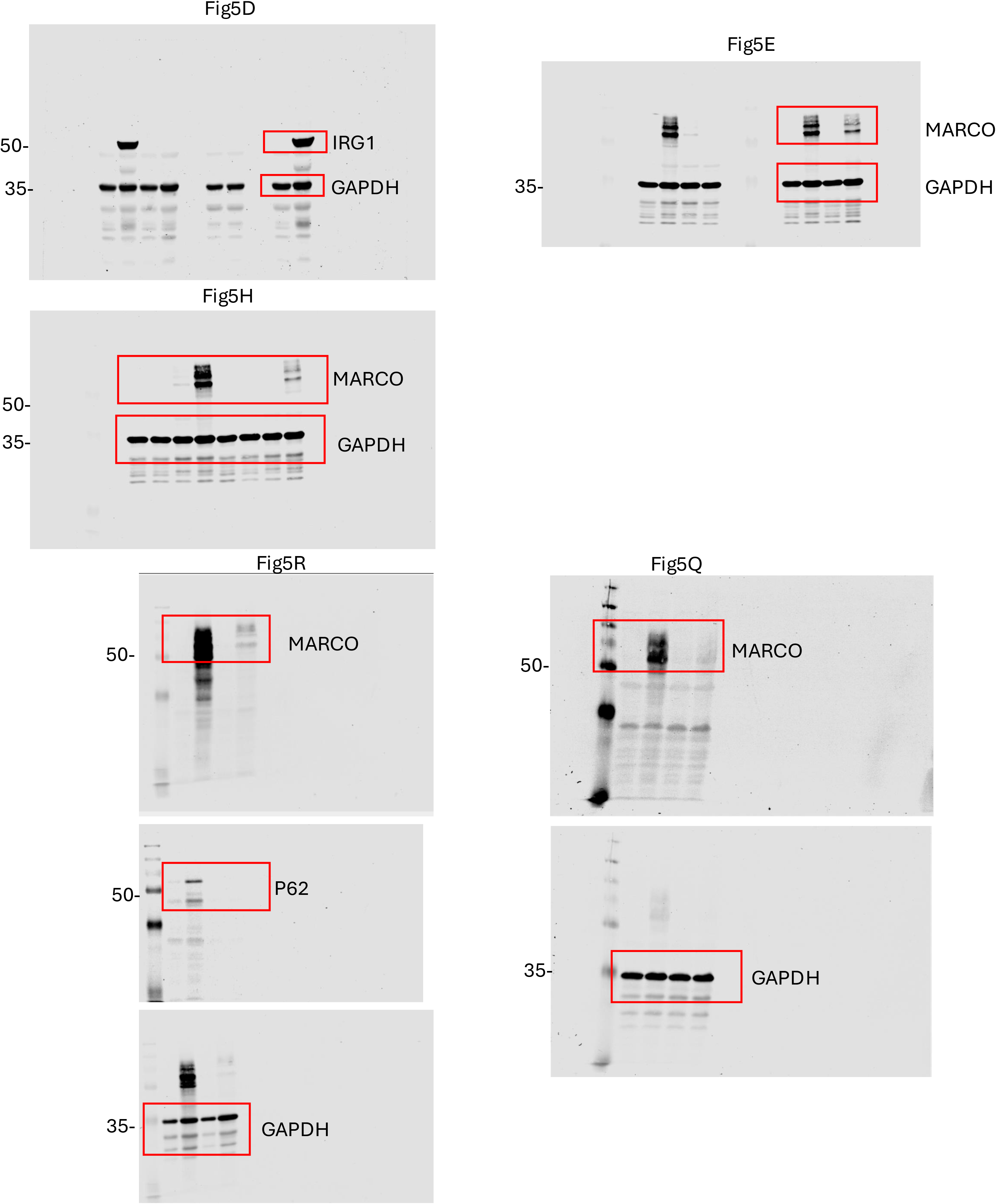
Uncropped western blots from Figure 5.

**Supplementary Figure 15:**
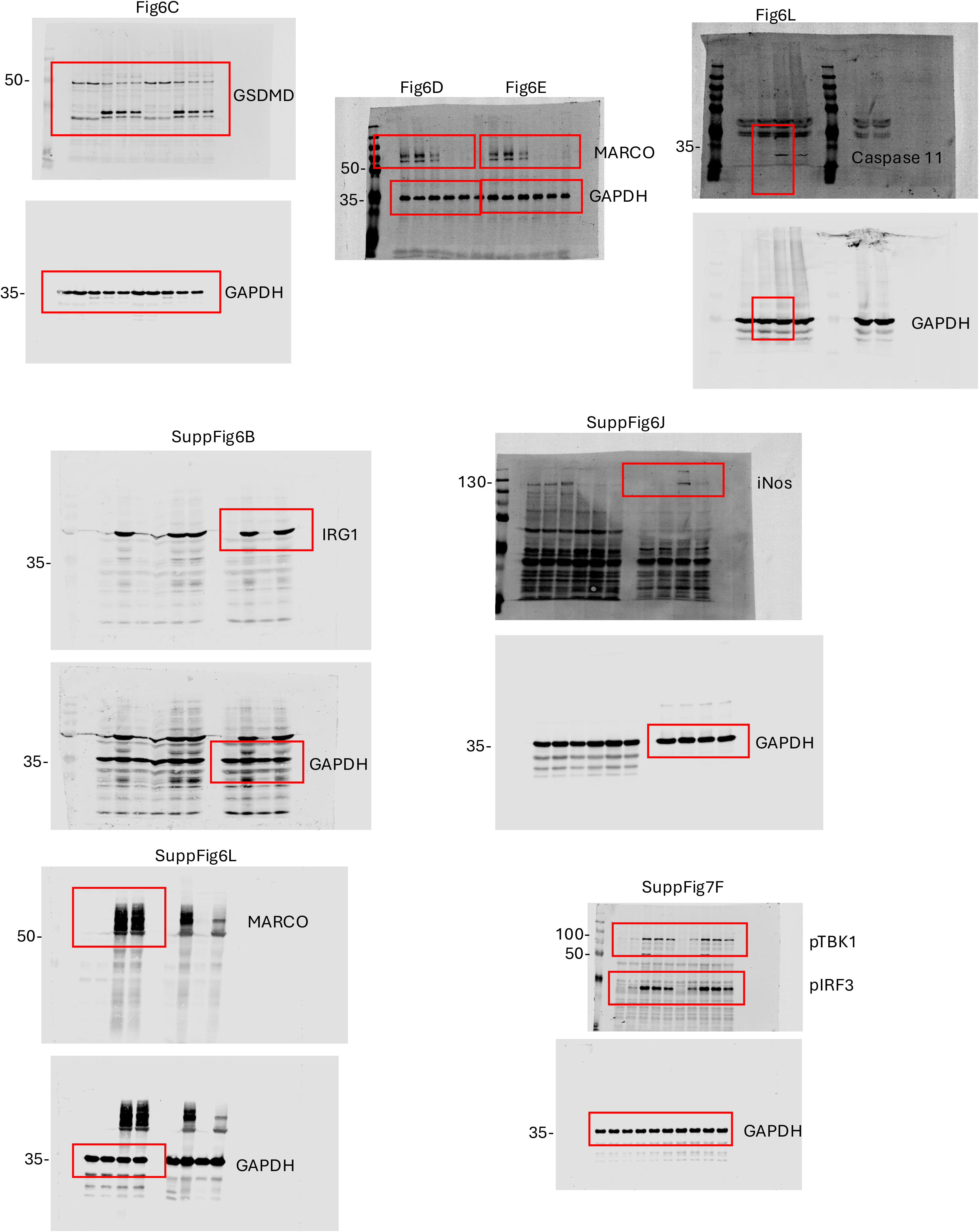
Uncropped western blots from Figure 6 and supp Figure 6 and 7.

